# Pioneer Transcription Factors Differentially Co-opt Bcl11a and Bcl11b to orchestrate Reprogramming

**DOI:** 10.64898/2025.12.19.694617

**Authors:** A. Trajkova, E. Zakiev, M. Mendoza-Ferri, I. Messahli, M. Ruel, I. Krossa, Y. Tapponnier, E. Dejonckere, E. Hunter, L. Claret, B. Martin, N. Gadot, F. Forest, C. Heinrich, N. Martin, D. Bernard, Y. Grinberg-Bleyer, M. Mendoza-Parra, A. Huyghe, F. Lavial

**Author notes:** Contacts for correspondence and. Jointly supervised (A. Huyghe, F. Lavial).

## Abstract

Pioneer transcription factors (TFs) orchestrate development, reprogramming, and cancer. Yet, the molecular mechanisms by which they cooperate with endogenous TFs and chromatin to trigger cell fate conversions remain largely unknown. Here, we identified antagonistic functions in reprogramming to pluripotency for the two paralogous somatic zinc finger TFs Bcl11a and Bcl11b. We reveal that Bcl11a and Bcl11b are initially co-expressed in mouse embryonic fibroblasts and then segregate in cellular intermediates respectively prone or refractory to reprogramming, transdifferentiation, and oncogenic transformation. They exert opposite functions - with Bcl11a promoting and Bcl11b hindering - the efficacy of induced pluripotent stem cells generation. During reprogramming, we uncover that Bcl11b safeguards cellular identity by persistently binding to differentiation genes with Runx1 in refractory intermediates. In contrast, in reprogramming intermediates, Bcl11a interacts with Oct4 and is initially displaced from MEF enhancers. Bcl11a then binds transiently and contributes to activate the E3 ubiquitin ligase Pja1 that regulates Smad3, thus promoting mesenchymal-epithelial transition and constraining senescence. Collectively, our work unveils how the differential repurposing of paralogous TFs by Oct4 orchestrates reprogramming to pluripotency.

## Introduction

Cell fate conversions (CFCs) occur in a wide range of normal and pathological processes including pluripotent reprogramming, transdifferentiation and oncogenic transformation^1, 2, 3^. Profound molecular changes contribute to the loss of cellular identity and to the gain of cellular plasticity during CFC^3, 4, 5, 6, 7, 8^. This is well illustrated by the multistep process of pluripotent reprogramming, triggered by Oct4, Sox2, Klf4 and cMyc (OSKM) to generate induced pluripotent stem (iPS) cells and by the combinatorial expression of three neural-lineage-specific transcription factors (Brn2, Ascl1 and Myt1l - BAM) that directly convert fibroblasts into induced neurons^3, 9^. Pluripotent reprogramming triggers massive changes of the epigenome, transcriptome but also of morphology and physiology, as exemplified by the switch of mouse embryonic fibroblasts (MEF) from a migratory mesenchymal to polarized epithelial state *via* mesenchymal-to-epithelial transition (MET)^4, 10, 11, 12, 13, 14, 15, 16, 17^. At the molecular level, the pioneer TFs OSK (i) redistribute somatic factors away from MEF enhancers to repress somatic identity and (ii) open pluripotent enhancers to activate the corresponding network^18, 19, 20^. Interestingly, OSKM also trigger poorly described epigenetic changes that generate cellular plasticity and promotes the emergence of alternative cell fates during reprogramming^12, 13, 14^. Partial loss of identity and gain of plasticity also contribute to tumor initiation and progression in certain contexts^20, 21^. Early transforming cells escape phenotypical restrictions and generate highly plastic cellular states, as shown recently in MEFs and in the lung^2, 4, 21, 22, 23^. Investigations into the interplays between reprogramming and oncogenic transformation have shown that oncogenic K-Ras (K-Ras^G12D^) promotes reprogramming in a context-dependent manner while a short induction of OSKM in pancreatic acinar cells promotes cancer formation^24, 25^. Recent comparative analysis of reprogramming and oncogenic transformation triggered in primary MEFs showed that both processes share an early molecular program^4, 26^. We thus speculated that dissecting the early stages of reprogramming, transdifferentiation and oncogenic transformation may uncover CFC regulators that could be targeted in regenerative biology. By focusing on early intermediate cells of the three CFC processes, we identified two critical regulators, namely the paralogous zinc finger TFs Bcl11a (B-cell lymphoma/leukemia 11a) and Bcl11b (B-cell lymphoma/leukemia 11b), that are initially co-expressed in MEFs. Upon induction of reprogramming, transdifferentiation or transformation, Bcl11a and Bcl11b segregate to become mainly expressed in cellular intermediates prone or refractory to CFC, respectively. They display opposite functions, Bcl11b acting as a roadblock, and Bcl11a a facilitator, during reprogramming, transdifferentiation and transformation *in vitro*. Mechanistically, during reprogramming, we unveiled that Bcl11b is mainly refractory to OSKM action and binds to differentiation genes with Runx1 in refractory intermediates. In contrast, Bcl11a is corrupted and converted into a pro-reprogramming factor by Oct4. Bcl11a is (1) displaced away from MEF enhancers by Oct4 to accelerate loss of cellular identity, (2) redistributed transiently to an enhancer region of the Pja1 locus to contribute to its activation that promotes MET and (3) binds pluripotent enhancers at late reprogramming stage.

## Results

### Identification of factors commonly regulating pluripotent reprogramming, neuron transdifferentiation and oncogenic transformation

To identify early regulators of reprogramming, transdifferentiation and transformation, we conducted single-cell RNA-sequencing (scRNA-seq) of primary MEFs induced to reprogram (*via* Oct4, Sox2, Klf4 and c-Myc (OSKM)), to transdifferentiate (*via* Brn2, Ascl1 and Myt1L (BAM)), or to transform (*via* mutated H-Ras and c-Myc combined with p53 depletion) for 3 days (Fig. 1a and Supplementary Fig. 1a)^1, 4, 27^. The number of cells per sample varied between 869 and 2633 cells for a total of 6585 cells. We next generated a diffusion map with the destiny package and obtained five different clusters using density-based clustering (HDBSCAN) (Fig. 1b-c). Three main early fates, corresponding to reprogramming (Cluster 5), transdifferentiating (Cluster 3), and transforming (Clusters 2 and 4) cells, emerged from the initial MEFs (Cluster 1). We recently demonstrated the emergence of cellular intermediates expressing *Bcl11b* and *Thy1* at low (double low DL cells) or high (double high DH cells) levels during reprogramming and oncogenic transformation^4^. In line with the scRNA-seq experimental design, we showed here that DL cells, FACS sorted at day3 but also at a later timepoint such as day7 of reprogramming or oncogenic transformation, are significantly more prone than DH cells to generate pluripotent or immortalized derivatives, as shown by Alkaline Phosphatase (AP) or immortalized foci staining, respectively (Supplementary Fig. 1b-e). Interestingly, cells downregulating *Bcl11b* and *Thy1* were found to emerge on the diffusion map during the three CFC processes (Fig. 1d-e and Supplementary Fig. 1f). EnrichR^15^ analysis revealed that DL cells harbor reduced levels of genes related to the EMT (*Tgfb1, Nnmt*), a crucial event for both reprogramming and cancer biology (Fig. 1f)^11,16^ and induced levels of genes related to the cell cycle (G2-M checkpoint) (*Ccnb2*, *Cenpa*), in line with changes in proliferation rate^17,18^. Bioinformatic analyses led us to identify transcripts homogeneously induced in DL cells, when compared with DH cells, that could constitute broad regulators of cell fate changes. Among the top 15 candidates, we noticed the presence of previously described regulators of reprogramming, such as Hmga1 and Hoxa10 (Fig. 1g and Supplementary Fig. 1g)^28, 29^. We also noticed the transcription factor Bcl11a being more abundant in DL cells (Fig. 1g-h). This finding was intriguing as we recently reported that its paralog Bcl11b acts as a broad-scope roadblock of CFC^4^. We therefore wondered whether Bcl11a also regulates CFC using a FACS-based strategy centered on the combined expression of Bcl11b and Thy1^4^. DH MEFs were FACS sorted to purity and Bcl11a knockdown (KD) was conducted prior to inducing reprogramming or transformation (Fig. 1i and Supplementary Fig. 1h-i). After 10 days of reprogramming or oncogenic transformation, in control settings, the DH cells were reduced to 5% and 19% of the whole population, respectively, because of their efficient conversion into alternative substates (Fig. 1i-j). Bcl11a depletion impaired the efficiency of such a conversion, leading to a significantly larger proportion of remaining DH cells (19% and 35% respectively) (Fig. 1i-j). Collectively, bioinformatic analyses of single cell transcriptomic data led to identify the Bcl11a TF as a facilitator of the early cellular changes in MEF.

**Fig. 1:**
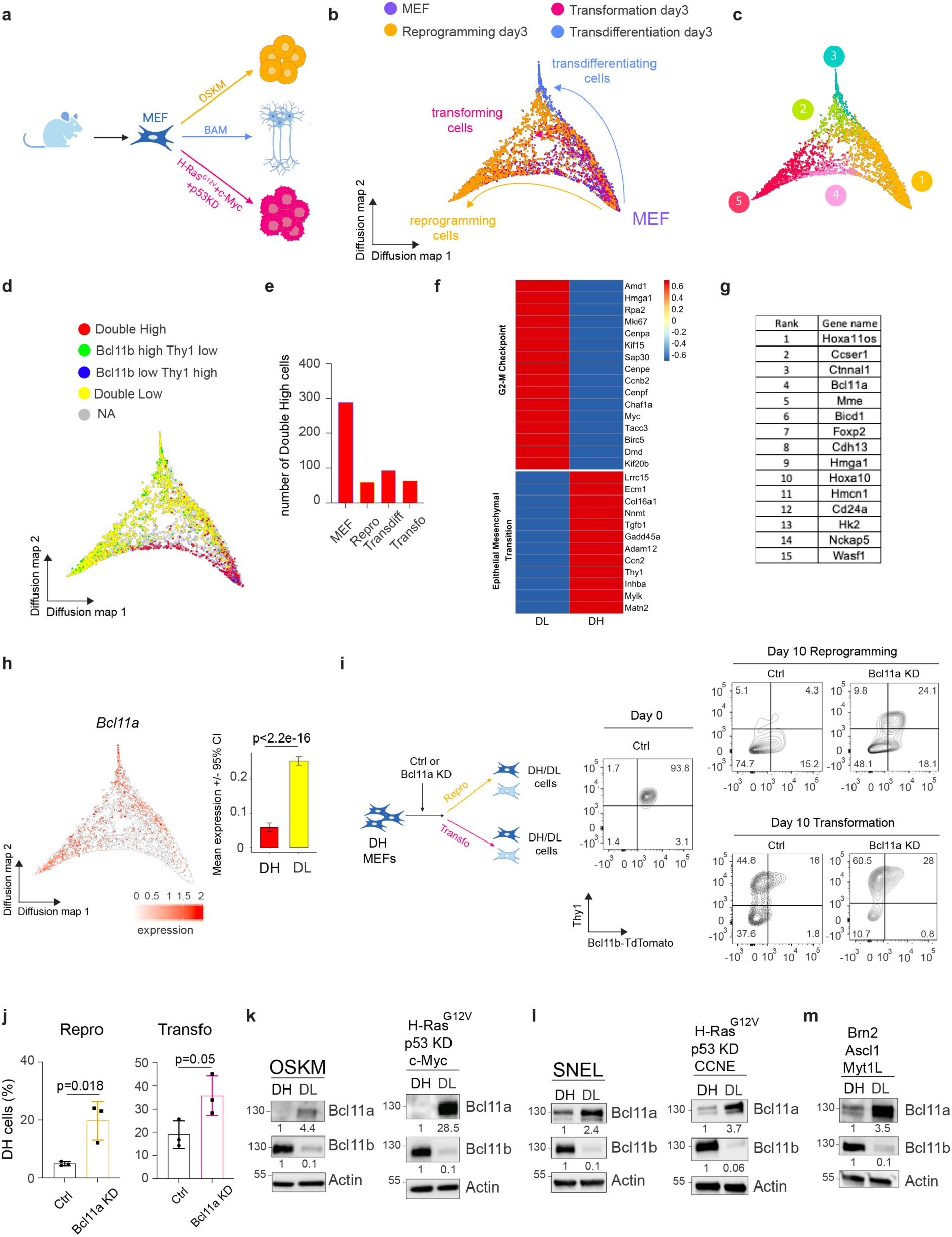
Distinct Bcl11a and Bcl11b expression pattern in cellular intermediates during MEF reprogramming, transdifferentiation and transformation. **a** Schematic diagram showing the strategy used to induce pluripotent reprogramming, neuron transdifferentiation or oncogenic transformation in MEFs. **b-c** Diffusion map stemming from scRNA-seq analysis of MEFs, reprogramming, transdifferentiating and transforming cells at day 3, where the color scales represent sample (**b)** or cluster affiliation (**c**). **d** Diffusion map based on the combined expression level of *Bcl11b* and *Thy1*. The thresholds for Low and High were 0.5 for Bcl11b and 1.5 for Thy1. **e** Bar chart depicting the number of double high cells in the different samples. **f** Heatmap of marker gene expression of the two major GO terms in DH and DL cells scaled by row. **g** List of the genes homogeneously induced in DL cells when compared with DH cells. **h** Expression profile of *Bcl11a* on the diffusion map (left panel) and a bar chart (right panel). **i** Left: schematic diagram of the experimental design. Right: FACS profile of Bcl11b (tdTomato) and Thy1 expression in MEF (day 0) and day 10 of reprogramming and transformation in Control (Ctrl) and Bcl11a KD condition. **j** Quantification of the percentage of Bcl11b/Thy1-expressing cells (DH) in Ctrl versus Bcl11a KD conditions during reprogramming (n=3 independent experiments, left panel) and transformation (n=3 independent experiments, right panel). The data represent means ± sd. **k** Western blot depicting Bcl11a, Bcl11b and Actin levels in DH and DL cells at day 5 of OSKM-mediated reprogramming (left panel) and HRas^G12V^, cMyc and shp53 induced oncogenic transformation (right panel). **l** Western blot depicting Bcl11a, Bcl11b and Actin expression in DH and DL cells at day 5 of reprogramming induced by Sall4, Nanog, Esrrb and Lin-28 (SNEL, left panel) and transforming induced by HRas^G12V^ and Cyclin E (CCNE) expression combined with p53 depletion (right panel). **m** Western blot depicting Bcl11a, Bcl11b and actin expression in DH and DL cells at day 5 of direct neuron conversion induced by Brn2, Ascl1 and Myt1L expression. Statistical significance was determined by Wilcoxon Ranks Sum test for h and two-tailed Student’s t-test for j.

#### Bcl11a expression segregates in cellular intermediates prone to CFC

We next monitored Bcl11a expression during reprogramming and oncogenic transformation. At the bulk level, Bcl11a is transiently upregulated at day2 for both processes before a progressive downregulation (Supplementary Fig. 1j). Among the reprogramming and transforming factors, the sole exogenous expression of Oct4 and c-Myc, as well as p53 depletion, is sufficient to trigger Bcl11a upregulation (Supplementary Fig. 1k). We next analyzed the dynamic expression of Bcl11a in cellular intermediates^4^. At day5 of reprogramming and oncogenic transformation, Bcl11a expression was mainly detected in DL cells, prone to CFC (Fig. 1k and Supplementary Fig. 1l). A high *Bcl11a* expression restricted to reprogramming intermediates was also observed during MEF and neutrophil reprogramming in published datasets (Supplementary Fig. 1m). To rule out the reliance of these results on the combination of factors we used, we also tested alternative sets, namely Sall4, Nanog, Esrrb and Lin28 (SNEL)^20^ for reprogramming and HRas^G12V^, Cyclin E^21^ and p53 depletion for transformation and observed similar results (Fig. 1l and Supplementary Fig. 1n). Bcl11a segregation in DL cells was also detected during MEFs transdifferentiation into induced neurons^3^ (Fig. 1m and Supplementary Fig. 1o). Collectively, we identified the segregation of the paralog TFs Bcl11a and Bcl11b as a broad hallmark of MEF-derived cellular intermediates respectively prone or refractory to CFC.

#### Bcl11a increases the efficiency of mouse and human pluripotent reprogramming *in vitro*

Given that Bcl11a is expressed at a high level in cellular intermediates prone to change identity, we studied its functional implication in reprogramming. Bcl11a depletion led to a 10-fold reduction in the number of AP+ iPSC colonies (Fig. 2a-c). We showed next that *Bcl11a* is expressed in human dermal fibroblasts (HDF) and transiently induced during reprogramming (Supplementary Fig. 2a)^30^. Bcl11a Crispr/CAS9-mediated depletion significantly reduces human iPS cells generation efficiency (Fig. 2d-f), indicating that Bcl11a promotes mouse and human reprogramming efficacy.

**Fig. 2:**
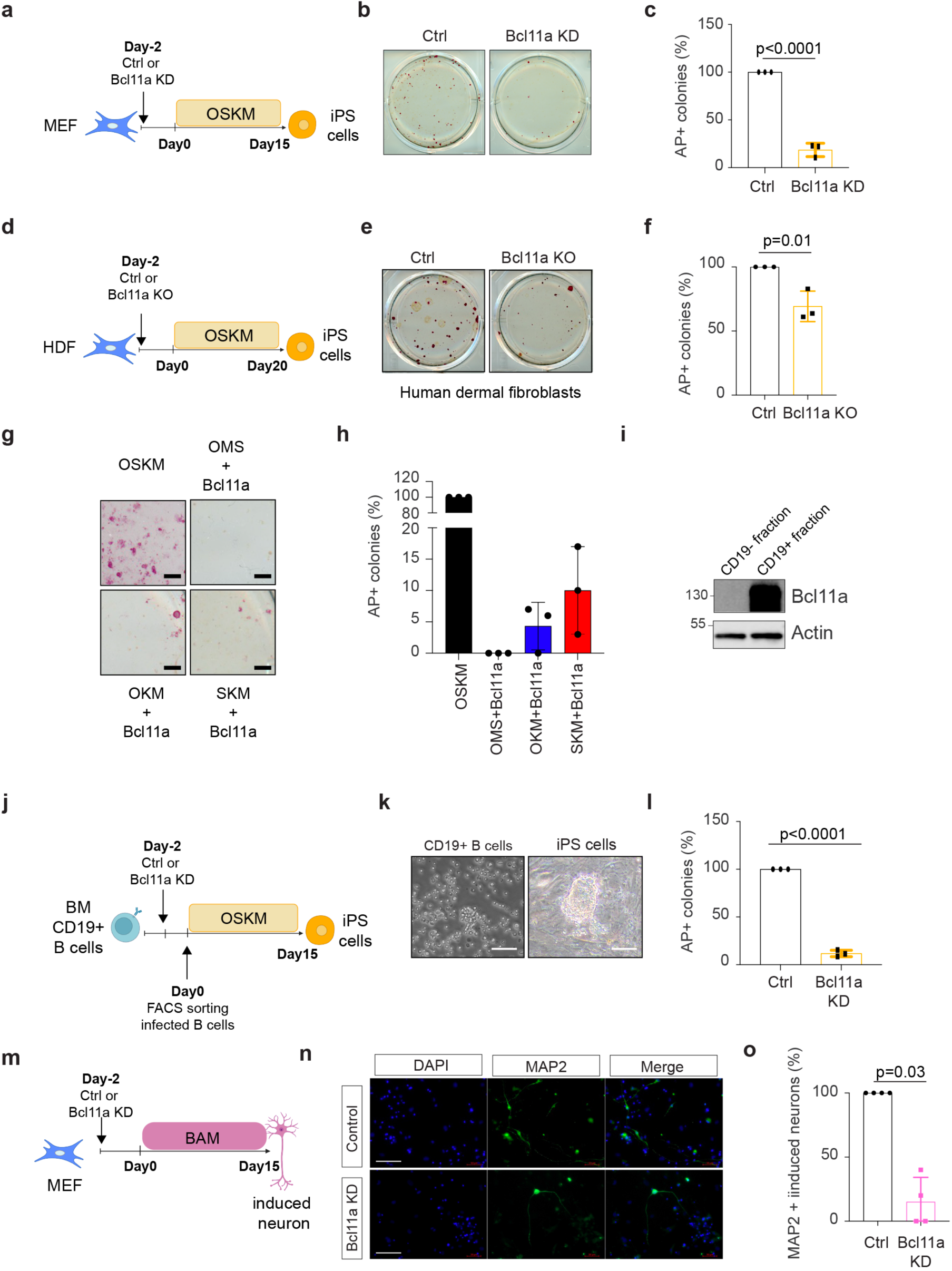
Bcl11a regulates the efficiency of reprogramming, transdifferentiation and transformation *in vitro*. **a** Schematic diagram of the experimental design to generate iPS cells from MEFs. **b** Representative images of iPS colonies stained for alkaline phosphatase (AP) activity. **c** Quantification of AP+ colonies (n=3 independent experiments). **d** Schematic diagram of the experimental design to generate iPS cells from HDFs. **e** Representative images of human iPS colonies stained for alkaline phosphatase (AP) activity. **f** Quantification of AP+ colonies (n=3 independent experiments). **g** Representative images of iPS colonies stained for alkaline phosphatase (AP) activity. Scale bar: 400 µm. **h** Quantification of AP+ colonies (n=3 independent experiments). **i** Western blot for Bcl11a and actin in CD19- and CD19+ cells. **j** Schematic of the experimental design of B cells reprogramming into iPSC. **k** Brightfield pictures of CD19+ B cells after bone marrow extraction (left panel) and iPS cells colony on inactivated OP-9 feeder following 15 days of reprogramming (right panel). Scale bar: 50µm. **l** Quantification of alkaline phosphatase positive (AP+) colonies (n=3 independent experiments). **m** Schematic diagram of the transdifferentiation of MEFs into induced neurons. **n** Immunofluorescence for MAP2. Scale bar: 100 µm. **o** Quantification of MAP2+ cells (n=4 independent experiments). **p** Schematic diagram of the experimental design of MEF oncogenic transformation. **q** Representative images of soft agar colonies. **r** Quantification of soft agar colonies. (n=3 independent experiments). The data represent means ± sd (**c, f, h, l, o, r**). Statistical significance was determined by two-tailed Student’s t-test (**c, f, h, l, o, r**).

We next showed that Bcl11a overexpression (OE) does not significantly improve reprogramming efficacy when combined with OSKM (Supplementary Fig. 2b-c). However, we asked if its action is capable of replacing some of the reprogramming factors. Among the reprogramming factors, c-Myc was found to be dispensable to generate mouse iPS cells^31^. We revealed that Bcl11a is able to generate AP+ iPS colonies in combination with OKM and SKM but not with OMS (Fig. 2g-h).

We next evaluated Bcl11a function during reprogramming of another somatic cell type. To that purpose, we extracted OSKM Dox-inducible Bone Marrow (BM) CD19+ B-lymphocytes highly expressing Bcl11a (Fig. 2i and Supplementary Fig. 2d). Reprogramming was induced with Dox treatment accompanied by coculture on OP-9 feeder cells, as previously reported^32^. Given the low transduction efficacy of CD19+ cells (<10% - Supplementary Fig. 2e), transduced cells were FACS sorted and replated at a similar density prior to Dox treatment (Fig. 2j). Bcl11a depletion led to a 10-fold decrease in the number of AP+ iPS colonies (Fig. 2k-l). Collectively, our data show that Bcl11a is not absolutely required but significantly increases the reprogramming efficiency of various mouse and human somatic cells.

#### Bcl11a increases the efficiency of direct neuron transdifferentiation and oncogenic transformation *in vitro*

To assess whether Bcl11a more broadly regulates another process of CFC, we focused on MEF transdifferentiation into induced neurons triggered by BAM, during which Bcl11a expression was high in DL cells (Fig. 1h, m and Supplementary Fig. 1o). Bcl11a depletion, prior to the induction of transdifferentiation, nearly abolished the emergence of MAP2+ induced neurons (Fig. 2m-o).

We finally assess whether Bcl11a also regulates the efficiency of oncogenic transformation *in vitro* as its expression is induced in DL cells (Fig. 1 k-l and Supplementary Fig. 1l, n). We found that Bcl11a depletion, prior to the induction of oncogenic transformation, led to a 3-fold decrease in the number of immortalized foci and 2-fold decrease in the number of transformed colonies in soft agar assays (Fig. 2p-r and Supplementary Fig. 2f-g). In contrast, Bcl11a OE had no significant effect in similar assays (Supplementary Fig. 2h-i). Collectively, our data indicate that Bcl11a is not absolutely required but promotes the efficiency of transdifferentiation and transformation *in vitro*.

#### Bcl11a and Bcl11b have different effects on MEF reprogramming trajectory

Given the antagonistic roles of Bcl11a and Bcl11b on MEF reprogramming, we wondered how they regulate cellular trajectories at the single cell resolution. ScRNA-seq was conducted on Ctrl, Bcl11a KD or Bcl11b KD cells at 6 different time points (Day0, Day3, Day6, Day9, Day12, Day14), corresponding to 3 reprogramming phases (Early day3-6; Mid day6-9 and Late day9-14) (Fig. 3a). After preprocessing 74,303 cells, HDBSCAN clustering on the dimensionality reduction embedding of uniform manifold approximation and projection (UMAP) led to the definition of 12 clusters of cells (Fig. 3b-c). We showed with a heatmap of differentially expressed genes distinct waves of transcriptomic changes occurring during reprogramming (Supplementary Fig. 3a). Previously published signatures, combined with marker genes and signatures from the PanglaoDB database^25^, led to the affiliation of clusters along the reprogramming of MEFs to iPS cells. Cluster 1 corresponded to MEFs while Clusters 5 and 9 reflect pluripotent cells expressing *Nanog* and *Zfp42* (Fig. 3d-e and Supplementary Fig. 3b-c)^13^. We next calculated reprogramming trajectories using the Slingshot R package (Fig. 3f). While a single trajectory led MEFs to reach the pluripotent clusters 5 and 9, two alternative trajectories led to the clusters 2 and 12, respectively (Fig. 3f). For Bcl11b KD cells, we observed a similar pattern of clustering to that of Ctrl cells (Fig. 3b, g). However, from Mid reprogramming, the percentage of Bcl11b KD cells reaching the pluripotent Clusters (5 and 9) was higher than Ctrl cells, consistent with the role of Bcl11b as a barrier to CFC (Fig. 3h). At the early stage, Bcl11a KD cells mostly followed Ctrl and Bcl11b KD cells (Fig. 3b) but diverged at Mid reprogramming where their trajectories led to non-pluripotent clusters, especially clusters 2, 3, 4, 8 and 12 (Fig. 3f-g, i and Supplementary Fig. 3d). Of note, certain cells from Cluster 12 started to activate the expression of the extra-embryonic markers *Gata4* and *Gata6* (Fig. 3j)^12, 13^ while cells from Cluster 4 expressed genes related to extracellular matrix and cell adhesion (Supplementary Fig. 3e). Moreover, as Bcl11a KD cells also fail to attain the pluripotent Clusters 5 and 9 (Fig. 3h), we wondered whether Bcl11a contributes to the activation of the endogenous Oct4 using Pou5f1-GFP reporter mice^26^. FACS analysis showed that the percentage of Pou5f1-GFP expressing cells was significantly decreased by Bcl11a depletion at both Mid and Late stages of reprogramming (Fig. 3k-l). Altogether, these results show that Bcl11a canalizes the reprogramming trajectory towards pluripotency and represses the expression of extra-embryonic markers.

**Fig. 3:**
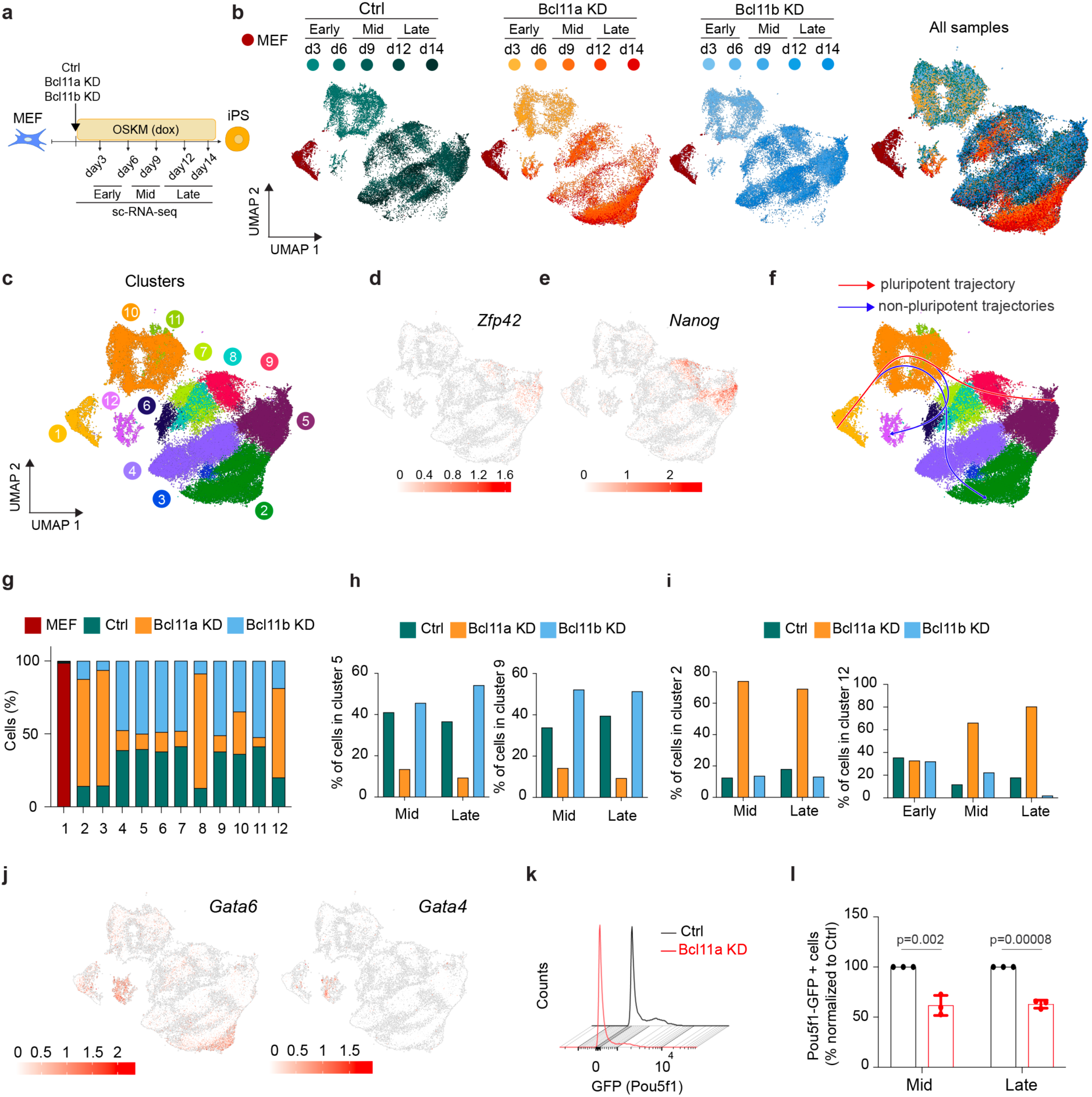
Depletion of Bcl11a and Bcl11b has different effects on reprogramming trajectories. **a** Schematic diagram of the experimental design for scRNA-seq of control (Ctrl), Bcl11a KD or Bcl11b KD reprogramming. **b** Uniform manifold approximation and projection (UMAP) plot for each condition (Ctrl, Bcl11a KD, Bcl11b KD) and merged settings. Colors indicate samples. **c** Merged UMAP with colors representing clusters. **d-e** Expression levels of *Zfp42* (**d**) and *Nanog* (**e**) on UMAP. **f** UMAP plot depicting the pseudo-time trajectories inferred with Slingshot. **g** Proportion of cells from each sample assigned to each cluster. **h** Proportions of cells assigned to Cluster 5 (left panel) and Cluster 9 (right panel) at the Mid and Late reprogramming stage as defined in (a). **i** Proportion of cells assigned to Cluster 2 (left panel) and Cluster 12 (right panel) at Early, Mid and Late of reprogramming. **j** Expression patterns of *Gata6* (left panel) and *Gata4* (right panel) on UMAP. **k** Representative FACS profile of Pou5f1-GFP positive cells in Ctrl and Bcl11a KD condition at late reprogramming stage. **l** Quantification of Pou5f1-GFP positive cells in Ctrl and Bcl11a KD settings at mid and late reprogramming stage (n=3 independent experiments). The data represent means ± sd. Statistical significance was determined by two-tailed Student’s t-test.

#### Bcl11a protects reprogramming cells from p16-mediated senescence and facilitates mesenchymal-to-epithelial transition (MET)

Next, we characterized cells within the non-pluripotent trajectories promoted by Bcl11a depletion, speculating that they may be affected by the established barriers of reprogramming, notably senescence or apoptosis^33, 34, 35^. Compared to Ctrl and Bcl11b KD cells, Bcl11a KD cells displayed a significantly higher senescence score during reprogramming, while apoptosis score was similar or even decreased (Fig. 4a and Supplementary Fig. 3f). Focusing on senescence, we assessed the expression of the known senescence inducers *p16 (Cdkn2a), p15 (Cdkn2b)* and *p21 (Cdkn1a)* in Cluster 2, and observed a striking increase in *p16* expression (Fig. 4b). Moreover, Bcl11a depletion triggered a significant induction of the senescence-associated transcripts *Ccl8* and *Cxcl2* at Late reprogramming (Supplementary Fig. 3g)^36^. We then devised a RNAi combinatorial strategy to understand the interplay between Bcl11a and p16. P16 KD was sufficient to partially revert Bcl11a KD phenotype for both induction of senescence transcripts (Fig. 4c and Supplementary Fig. 3h) and reprogramming efficacy (Fig. 4d-e). Thus, Bcl11a prevents the induction of p16-mediated senescence during MEF reprogramming.

**Fig. 4:**
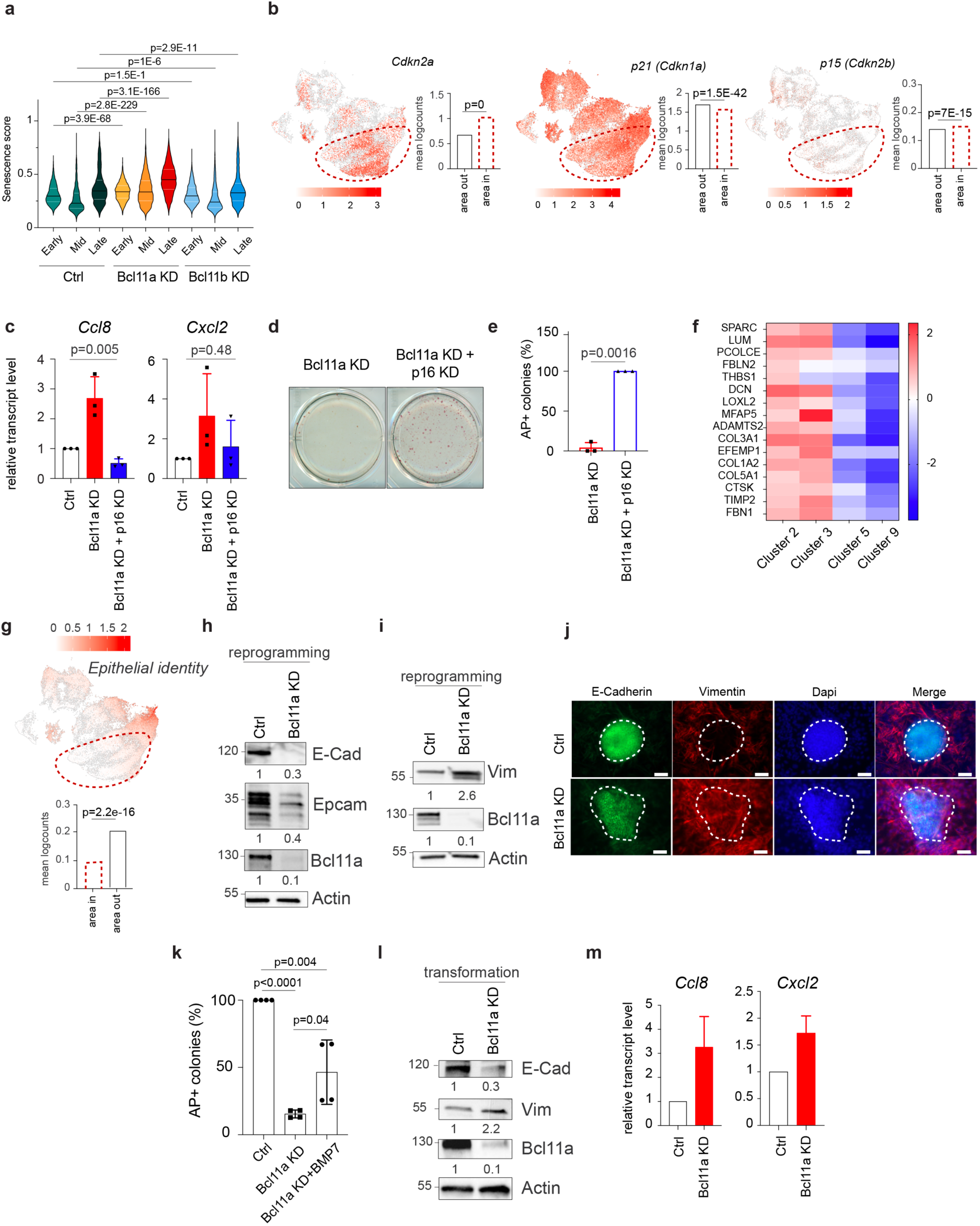
Bcl11a constrains senescence and promotes MET during reprogramming. **a** Violin plot depicting the level of a senescence signature score (from Ref.^13^). **b** UMAP plot for *Cdkn2a* (left), *Cdkn1a* (middle) and *Cdkn2b* (right) expression level in all cells (Ctrl, Bcl11a and Bcl11b KD). **c** Graph depicting *Ccl8* and *Cxcl2* relative expression levels in Ctrl, Bcl11a KD and Bcl11a/p16 KD condition at late reprogramming stage (n=3 independent experiments). **d** Representative images of iPSC colonies stained for alkaline phosphatase (AP) activity. **e** Quantification of AP+ colonies (n=3 independent experiments). **f** Heatmap depicting expression levels of genes related to the Extracellular Matrix Organization GO term according to Reactome 2022 signature in Cluster 2, 3, 9 and 5. **g** UMAP plot depicting the expression level of an Epithelial identity signature (from Ref. ^13^). **h-i** Western blot for E-Cadherin, Epcam, Bcl11a, Vimentin and Actin in Ctrl and Bcl11a KD settings at late reprogramming stage. **j** Immunofluorescence for E-cadherin and Vimentin. Scale bar: 50 µm. **k** Quantification of AP+ colonies after 15 days of reprogramming in Ctrl, Bcl11a KD and Bcl11a KD + BMP7 settings. Cells were treated with recombinant BMP7 for the first 5 days of reprogramming (2nM) (n=4 independent experiments). **l** Western blot for E-Cadherin, Vimentin, Bcl11A and Actin in Ctrl and Bcl11a KD settings at late transforming stage. **m** Graph depicting *Ccl8* and *Cxcl2* relative expression levels in Ctrl and Bcl11a KD condition at day4 of oncogenic transformation (n=two independent experiments). The data represent means ± sd (**c, e, k, m**). Statistical significance was determined by two-tailed Student’s t-test (**c, e, k**) and Wilcoxon Ranks Sum test (**a, b, g**).

In addition, Clusters 2 and 3, which are mainly composed of Bcl11a-KD cells, were enriched for genes related to the extracellular matrix (ECM) (Supplementary Fig. 3i-j) and mesenchymal identity (*Col1a2*, *Col5a1*, *Col3a1, Mfap5* and *Sparc)* compared to the pluripotent Clusters 5 and 9 (Fig. 4f)^37^. As MEF reprogramming is accompanied by an early MET, we wondered whether Bcl11a regulates the process. Not only did cells in Clusters 2 and 3 display a low epithelial identity score (Fig. 4g), they also failed to induce the epithelial markers E-Cadherin and Epcam (Fig. 4h) and maintained a high expression of the mesenchymal marker Vimentin at Late reprogramming (Fig. 4i). In line with it, while nascent control iPS colonies were tight and dome-shaped in Vimentin-low areas, emerging Bcl11a KD iPS colonies harbored loosed morphology in Vimentin-expressing areas (Fig. 4j). Finally, we showed that the Bcl11a KD phenotype was partially reverted by inducing MET using recombinant BMP7 treatment (Fig. 4k). Interestingly, Bcl11a was found to promote MET (Fig. 4l) and constrain the induction of the SASP molecules *Ccl8* and *Cxcl2* (Fig. 4m) during oncogenic transformation *in vitro*, as shown by western blot and Q-RTPCR, respectively.

These results indicate that Bcl11a prevents p16-mediated senescence and favors MET during reprogramming.

#### Bcl11a and Bcl11b interactomes are poorly overlapping in MEFs

We next attempted to further explore the antagonistic functions of Bcl11a and Bcl11b during CFC. Protein sequence alignment analysis indicated a 58,6% identity while domain analysis using PFAM showed the presence of C2H2 Zinc finger domains in both paralogs (Supplementary Fig. 4a). We next compiled Bcl11a and Bcl11b interactomes by performing Immuno-Precipitation coupled with Mass Spectrometry (IP-MS) in MEFs. We identified 239 interactants for Bcl11a and 278 for Bcl11b (Fig. 5a). Among those, and in line with previously published studies, chromatin-interacting complexes such as SWI/SNF (Smarcc2) and NuRD (Chd1) associated with Bcl11a^38, 39^. Unexpectedly, we unveiled an association with Smad proteins (Smad2, 3 and 5), known to be involved in BMP and TGFβ signaling^40^. Bcl11b interactants encompassed nucleosome assembly and transcription-involved proteins (Nap1l1, Pura and Dxd1). Overall, Bcl11a and Bcl11b shared less than 10% interactants (30), indicating that both TFs have mainly distinct interactomes in MEFs despite a certain degree of sequence similarity.

**Fig. 5:**
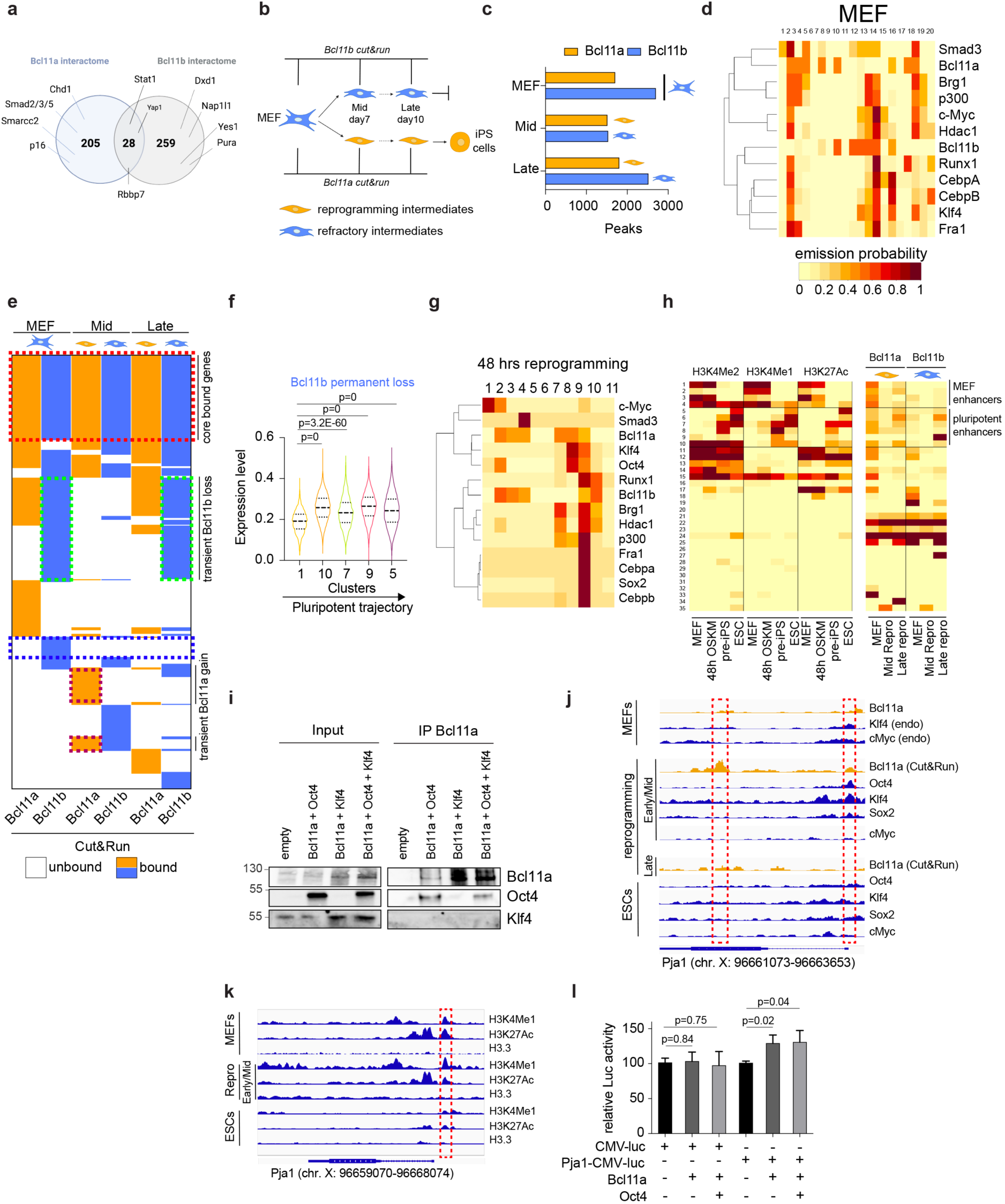
Bcl11a and Bcl11b distribution on the chromatin during reprogramming. **a** Venn diagram of Bcl11a and Bcl11b interactants in MEFs identified by IP-MS. **b** Schematic of the experimental design for Cut&Run analysis in MEF, reprogramming and refractory intermediates at Mid and Late reprogramming. **c** Number of binding peaks for Bcl11a and Bcl11b. **d** Chromatin state analysis comparing Bcl11a and Bcl11b binding sites in MEFs with ChIP-seq datasets in MEFs from ref.^7^. **e** Clustering of Bcl11a and Bcl11b bound genes in MEF together with Bcl11a binding in reprogramming intermediates and Bcl11b in refractory intermediates at Mid and Late stage of reprogramming. **f** Graph depicts the mean expression level of the 23 Bcl11b-bound genes losing permanently binding during reprogramming. **g** Chromatin state analysis comparing Bcl11a and Bcl11b binding sites at the Mid stage of reprogramming with ChIP-seq datasets in MEFs exposed to OSKM expression for 48 hours from ref.^7^. **h** ChromHMM analysis comparing chromatin states with Bcl11a and Bcl11b binding. **I** Co-immunoprecipitation of Bcl11a with Oct4 and Klf4 in 293T cells. **j-k** Snapshot of indicated genomics data at the Pja1 genomic locus. **l** Luciferase assay on 293T cells showing the binding of Bcl11a and Oct4 on the Pja1 enhancer. The data represent means ± sd (**l**). Statistical significance was determined by two-tailed Student’s t-test (**l**) or Wilcoxon Ranks Sum test (**f**).

#### Bcl11a and Bcl11b chromatin binding features are different in MEFs

We then explored the distribution of Bcl11a and Bcl11b on the chromatin by conducting Cut&Run assays in MEFs but also in FACS-sorted reprogramming intermediates (mainly expressing Bcl11a) and refractory intermediates (mainly expressing Bcl11b) at two timepoints (Mid and Late) of reprogramming (Fig. 5b).

In MEFs, Bcl11a binds to 227 genes (1,701 sites) and Bcl11b to 261 genes (2,695 sites), with 160 genes in common, as exemplified by the Tead1 locus (Fig. 5c and Supplementary Fig. 4b-c). More globally, the distribution of Bcl11a and Bcl11b peaks in regulatory elements appeared relatively similar (Supplementary Fig. 4d). DREME motif analysis led to predict Bcl11a and Bcl11b binding motifs. Of note, Tomtom comparison of the Bcl11a motif with the highest p-value indicated similarity with Mafk and Nfe2l1 motifs, consistent with previous works (Supplementary Fig. 4e)^41, 42^, but also unexpectedly Smads (Supplementary Fig. 4f), in line with our IP-MS results. To characterize the chromatin environment at sites engaged by Bcl11a and Bcl11b, we next built a chromatin state model with 20 states using ChromHMM and datasets of somatic TFs genome-wide binding in MEF^7, 43^. Bcl11a was preferentially co-occupying chromatin sites with Smad3 while Bcl11b was preferentially associated with Runx1, as recently reported in immune cells (Fig. 5d)^4, 44, 45, 46^. Altogether these results highlight that Bcl11a and Bcl11b preferentially associate with different somatic TFs on MEF chromatin.

#### Bcl11b remains closely associated with Runx1 in refractory intermediates

Having explored the preferential binding features of Bcl11a and Bcl11b to chromatin in MEFs, we next wondered how this binding evolved during reprogramming. The overall number of Bcl11a and Bcl11b peaks was not severely impacted by OSKM expression (Fig. 5c). We then clustered the Bcl11a and Bcl11b bound genes based on their dynamic behavior during reprogramming. We noticed first that more than half of the genes commonly bound by Bcl11a/Bcl11b in MEFs were also bound by either Bcl11a or Bcl11b in intermediates (143/160 (89%) at Mid and 94/160 (58%) at Late stage) (Fig. 5e – red dashed box). This core set of genes, related to housekeeping functions, remains expressed at similar level throughout reprogramming (Supplementary Fig. 4g). We next focused on the dynamic loss and gain of Bcl11a and Bcl11b binding during reprogramming (Fig. 5e). In refractory intermediates mainly expressing Bcl11b, 57% (150/261) of the genes occupied by Bcl11b in MEFs lost transiently or permanently its binding during reprogramming (Fig. 5e). Among those, the permanent loss of Bcl11b binding was correlated with a rapid and permanent upregulation of the corresponding genes during reprogramming, suggesting that Bcl11b acts as a transcriptional repressor (Fig. 5e – blue dotted box and 5f). Transient Bcl11b loss occurs on a very large proportion of genes (74% - 112/150) related to differentiation (Acer1, Ptpn2, Fstl5, Edrf1, Lage3 and Sema6c) or pluripotency (Pcbp1)^47^, suggesting that Bcl11b may thus be refractory to OSKM-mediated redistribution in refractory intermediates (Fig. 5e – green dashed boxes). Consistently, analysis of chromatin trajectories during reprogramming indicated that Bcl11b remained more closely bound with Runx1 than with reprogramming factors (Fig. 5g)^7^. We conclude here that Bcl11b remains associated with Runx1 and is largely refractory to OSKM-mediated redistribution.

#### Bcl11a binding to the MEF enhancer landscape is redirected by Oct4 in reprogramming intermediates

In reprogramming intermediates, Bcl11a was driven away from 50% (114/227) of the genes that it initially occupied in MEFs. Chromatin state analysis revealed that, in contrast to its paralog, Bcl11a binding was strongly influenced by OSKM and was particularly associated with Oct4 and Klf4 at early reprogramming stage (Fig. 5g). As Oct4 and Klf4 were found to alter the enhancer landscape during reprogramming, we evaluated Bcl11a and Bcl11b binding on MEF and pluripotency enhancers, as defined previously using H3K4me2, H3K4me1 and H3K27ac histone marking in MEF and embryonic stem cells (Fig. 5h)^7^. Strikingly, Bcl11a was associated with both MEF- (chromatin trajectories 1-4) and pluripotent- (chromatin trajectories 5-10) enhancers in MEF, suggesting that it contributes to sustain cellular identity. Moreover, this binding was strongly reduced in reprogramming intermediates (Fig. 5h). In line with this, Bcl11a was found to interact with Oct4 during reprogramming using co-immunoprecipitation assays in 293 T cells (Fig. 5i) but also by IP-MS conduced on MEFs induced to reprogram for 5 days (Supplementary Table 1). Therefore, we conclude that Bcl11a binding to the MEF enhancer landscape is redirected by the reprogramming factor Oct4 in reprogramming intermediates.

#### Bcl11a regulates Pja1 and Smad3 levels during reprogramming

To explain the role of Bcl11a as a facilitator of reprogramming, we hypothesized that Oct4 might redistribute and corrupt it to regulate key genes. We focused for this reason on the 99 genes that acquired Bcl11a binding in reprogramming intermediates. Among those, 48 genes were transiently bound by Bcl11a and upregulated early during reprogramming, suggesting that Bcl11a might be critical to ensure the proper activation of those genes (Fig. 5e – purple dashed boxes and Supplementary Fig. 4h). Within these genes, we identified ubiquitination-associated genes (Pja1, Ube2g2, Eif3h, Ubtd1) and showed in particular that the E3 ubiquitin ligase Pja1 locus, described to suppress TGF-β signaling by regulating Smad2/3, is also transiently bound by Oct4 and Klf4 during reprogramming (Fig. 5j)^48^. Of note, the newly bound region is located next to an active enhancer decorated by H3K4me1 and H3K27ac in MEFs and during reprogramming (Fig. 5k). By cloning the region bound by Bcl11a and Oct4/Klf4 next to the Luciferase gene, we demonstrated that it behaves as a regulatory region used by the two transcription factors to increase Pja1 transcription (Fig. 5l). In line with it, Bcl11a KD was found to reduce Pja1, and increase Smad3 level, during reprogramming (Supplementary Fig. 4i-j). Bcl11a therefore regulates Pja1 and Smad3 level during reprogramming.

#### Bcl11a-mediated induction of Pja1 promotes reprogramming

We next assessed whether Bcl11a-mediated induction of Pja1 and Smad3 regulates reprogramming. First, we found that Pja1 depletion reduces significantly reprogramming efficiency (Fig. 6a-b and Supplementary Fig. 4k-l). As expected, both Bcl11a and Pja1 depletion were found to increase Smad3 level and to constrain the emergence of epithelial cells, as shown by western blot for E-Cadherin and Epcam (Fig. 6c). However, while Smad3 depletion significantly reverted the induction of the senescence associated transcripts *Cxcl1*, *Cxcl2* and *CCl8* triggered by Bcl11a KD (Fig. 6d and Supplementary Fig. 4m), it did not restore reprogramming efficiency, indicating Smad3-independent functions for Bcl11a (Fig. 6e-f). In contrast, Pja1 exogenous expression was found to significantly rescue the reduction of reprogramming efficiency triggered by Bcl11a depletion (Fig. 6g-h). Hence, we propose that Bcl11a-mediated induction of Pja1 regulates reprogramming (Fig. 6i).

**Fig. 6:**
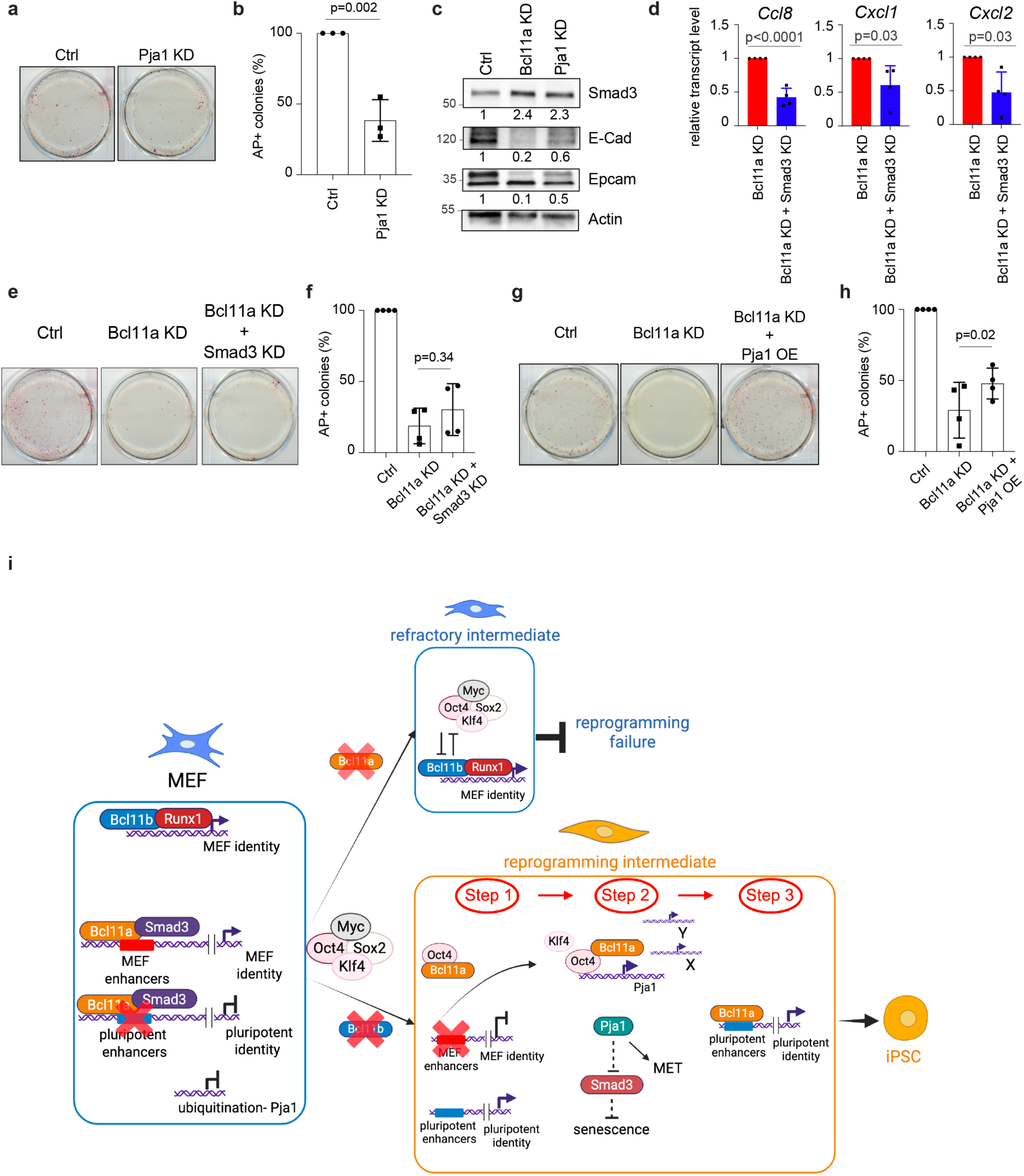
Bcl11a-mediated induction of Pja1 promotes reprogramming. **a** Representative images of iPSC colonies stained for alkaline phosphatase (AP) activity upon Pja1 KD. **b** Quantification of AP+ colonies (n=3 independent experiments). **c** Western blot of Smad3, E-cadherin and Epcam levels in Ctrl, Bcl11a KD and Pja1 KD settings at late stage of reprogramming. **d** Graph depicting *Ccl8, Cxcl1* and *Cxcl2* relative expression levels in Bcl11a KD and Bcl11a/Smad3 KD condition at late reprogramming stage (n=4 independent experiments). **e** Representative images of iPSC colonies stained for alkaline phosphatase (AP) activity upon Bcl11a KD and Bcl11a/Smad3 KD. **f** Quantification of AP+ colonies (n=4 independent experiments). **g** Representative images of iPSC colonies stained for alkaline phosphatase (AP) activity upon Bcl11a KD and Bcl11a/Pja1 KD. **h** Quantification of AP+ colonies (n=4 independent experiments). **i** Model. Bcl11a and Bcl11b are differentially co-opted by pioneer TF during cell fate conversion. In MEF, the distribution and the molecular partners of Bcl11a and Bcl11b are different. Bcl11a is particularly associated with the enhancer landscape. In refractory intermediates, Bcl11b remains associated with Runx1 to safeguard cellular identity and oppose OSKM-mediated redistribution. In reprogramming intermediates, Bcl11a binding to MEF- and pluripotency-enhancers is diminished and rather associates with Oct4 (step 1). Oct4, Klf4 and Bcl11a transiently bound to the Pja1 locus to ensure its activation that leads to TGF-β dampening via Smad3 and MET promotion (step 2). At the late stage of reprogramming, Bcl11a is bound to pluripotent enhancers (step 3). The data represent means ± sd (**b, d, f, h**). Statistical significance was determined by two-tailed Student’s t-test (**b, d, f, h**).

## Discussion

In this work, we identified the paralog TFs Bcl11a and Bcl11b as antagonistic regulators of pluripotent reprogramming. The comparison of the transcriptomic changes occurring in individual reprogramming, transdifferentiating and transforming cells pinpoints biological processes related to the reorganization of ECM and the EMT process but also candidate regulators such as Bcl11a and Bcl11b. Of note, Bcl11a and Bcl11b were previously described in immune and neuronal cells to trigger differentiation into distinct fates by compensating/repressing each other^49, 50^. In the present work, we revealed that Bcl11a and Bcl11b are co-expressed in MEFs and segregate respectively in cellular intermediates respectively prone or refractory to reprogramming, oncogenic transformation and transdifferentiation into induced neurons. This molecular switch appears however to be restricted to MEFs *in vitro*, as both factors are not expressed during pancreatic cell reprogramming *in vivo* (Supplementary Fig. 4n). As Bcl11a was found to promote the efficiency of a variety of processes, including reprogramming of mouse fibroblasts, mouse B cells and human dermal fibroblasts as well as the transdifferentiation of MEFs into neurons, it emerges as a broad facilitator of cell fate conversions. It appears however that Bcl11a plays a more limited role in oncogenic transformation.

By inferring the pseudo-time trajectories of reprogramming of control and Bcl11a- and Bcl11b-depleted cells, we showed that Bcl11a promotes reprogramming by (i) fostering MET, (ii) limiting p16-mediated senescence and (iii) constraining extra-embryonic gene expression. It will be interesting to further investigate whether the DNA repair function of Bcl11a, recently described in triple negative breast cancer, is required to canalize cellular outcomes during reprogramming^51^.

By combining IP-MS with Cut&Run approaches, we showed that Bcl11a and Bcl11b present common but also specific partners/target genes. In line with this, chromatin state analysis compiling our dataset with published ChIP-seq for other somatic TFs and histone marks^7, 43^ during reprogramming indicated that Bcl11a is bound on the MEF enhancer landscape on both MEF- and pluripotent-enhancer, in striking contrast with Bcl11b. Because Oct4 binds the enhancer landscape at the early stage of reprogramming and interacts with Bcl11a^7^, we speculate that the differences of epigenetic landscape occupancy between the two paralog TFs might be responsible for the differential co-optation of the somatic TFs by OSKM, even if additional investigations will be required. Indeed, striking differences in the behavior of the two TFs were identified after OSKM induction. In refractory intermediates, despite a certain degree of redistribution, Bcl11b remains co-bound with Runx1 on differentiation genes to secure MEF identity, behaving as expected as a somatic gatekeeper^52^. In contrast, Bcl11a is repurposed by Oct4. The redistribution of somatic TFs by reprogramming factors was proposed to contribute to silence MEF- and activate pluripotent-enhancer^7^. Here, we rather propose a tripartite model in which the somatic TF Bcl11a (1) is first displaced from MEF enhancers to accelerate loss of cellular identity, (2) then binds together with Oct4 the Pja1 locus to enhance its transcription to promote MET^41,46^ and (3) is finally bound to pluripotent enhancers at late stage of reprogramming (Fig. 6i). Bcl11a behaves therefore as a highly versatile TF co-opted by Oct4 to acquire pro-reprogramming functions, unlike Bcl11b. These findings expand the recent concept of Molecular Versatility^53^ that we illustrated using pluripotency progression as a paradigm to alternative processes such as reprogramming.

Finally, we identified multilayered interplay between Bcl11a and Smad3. Bcl11a and Smad3 interact together and co-bind on the chromatin in MEFs (Fig. 5a, d) and Bcl11a controls indirectly Smad3 level (Fig. 6c). As Smad3 level was proposed as a broad-scope potentiator of CFC^43^, it remains to be investigated (i) whether similar Pja1-mediated regulation of protein stability occurs during oncogenic transformation and neuron transdifferentiation and (ii) whether it involves Smad3 and/or alternative substrates.

By dissecting how paralogous somatic TFs can be differentially co-opted to acquire highly divergent functions *via* the forced expression of pioneer TFs, our work might offer new perspectives for advancing regenerative medicine and developing targeted cancer therapies.

---

**Supplementary Fig. 1:**
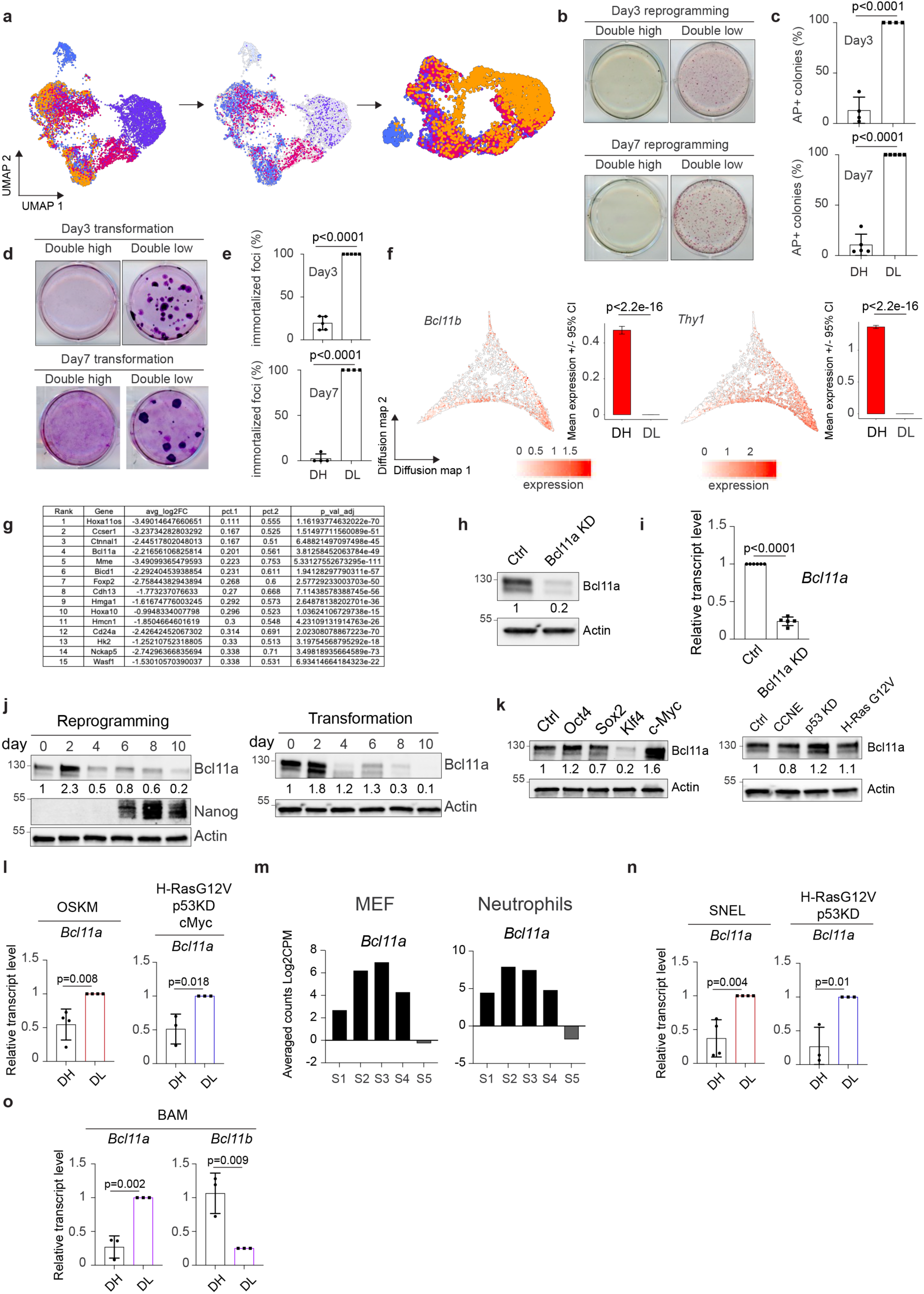
Distinct Bcl11a and Bcl11b expression in cellular intermediates during CFC in MEFs. **a** UMAP depicting the removal of the uninfected cells. Total cells are shown on the left panel, uninfected cells are presented on the middle panel (with no expression of the reprogramming, transdifferentiation or transformation factors) and the new UMAP generated on the restricted cells is presented on the right panel. **b** Representative images of iPSC colonies stained for alkaline phosphatase (AP) activity after 3 days (upper panel) or 7 days (lower panel) of Bcl11b/Thy1 double low and double high cell sorting. **c** Quantification of AP+ colonies (n=4/5 independent experiments). **d** Representative images of immortalized foci after 3 days (upper panel) or 7 days (lower panel) of Bcl11b/Thy1 double low and double high cell sorting. **e** Quantification of foci (n=4/5 independent experiments). **f** UMAP of the expression profile of *Bcl11b* and *Thy1* with quantitative bar chart. **g** List of the genes homogeneously induced in DL cells when compared with DH cells. **h** Western Blot for Bcl11a and Actin in Ctrl and Bcl11a KD MEF. **i** RT-qPCR analysis of *Bcl11a* relative expression level in Ctrl and Bcl11a KD MEF (n=5 independent experiments). **j** Western blot for Bcl11a, Nanog and Actin during reprogramming (left panel) and for Bcl11a and Actin during oncogenic transformation (right panel) at day 0, 2, 4, 6, 8 and 10. **k** Western blot for Bcl11a and Actin in the indicated settings of overexpression in MEFs. **l** RT-qPCR analysis of *Bcl11a* relative expression levels in prone and refractory intermediates at day 5 of OSKM-mediated reprogramming (left) (n=4 independent experiments) and of HRasG12V, cMyc, shp53 triggered oncogenic transformation (right) (n=3 independent experiments). **m** RNA-seq data from ref.^54^ showing *Bcl11a* transient upregulation during defined steps (S1 to S5) of fibroblast and neutrophil reprogramming into iPS cells. **n-o** RT-qPCR analysis of *Bcl11a* and *Bcl11b* levels in prone and refractory intermediates at day 5 of pluripotent reprogramming and oncogenic transformation induced by alternative reprogramming (n=4 independent experiments) and transforming cocktails (n=3 independent experiments) (**n)** and of BAM induced transdifferentiation (n=3 independent experiments) (**o**). The data represent means ± sd (**c**, **e**, **i**, **l**, **n**, **o**). For statistical significance two-tailed student’s T-test (**c, e, i, l, n, o**) and Wilcoxon Ranks Sum (**f**) were used.

**Supplementary Fig. 2:**
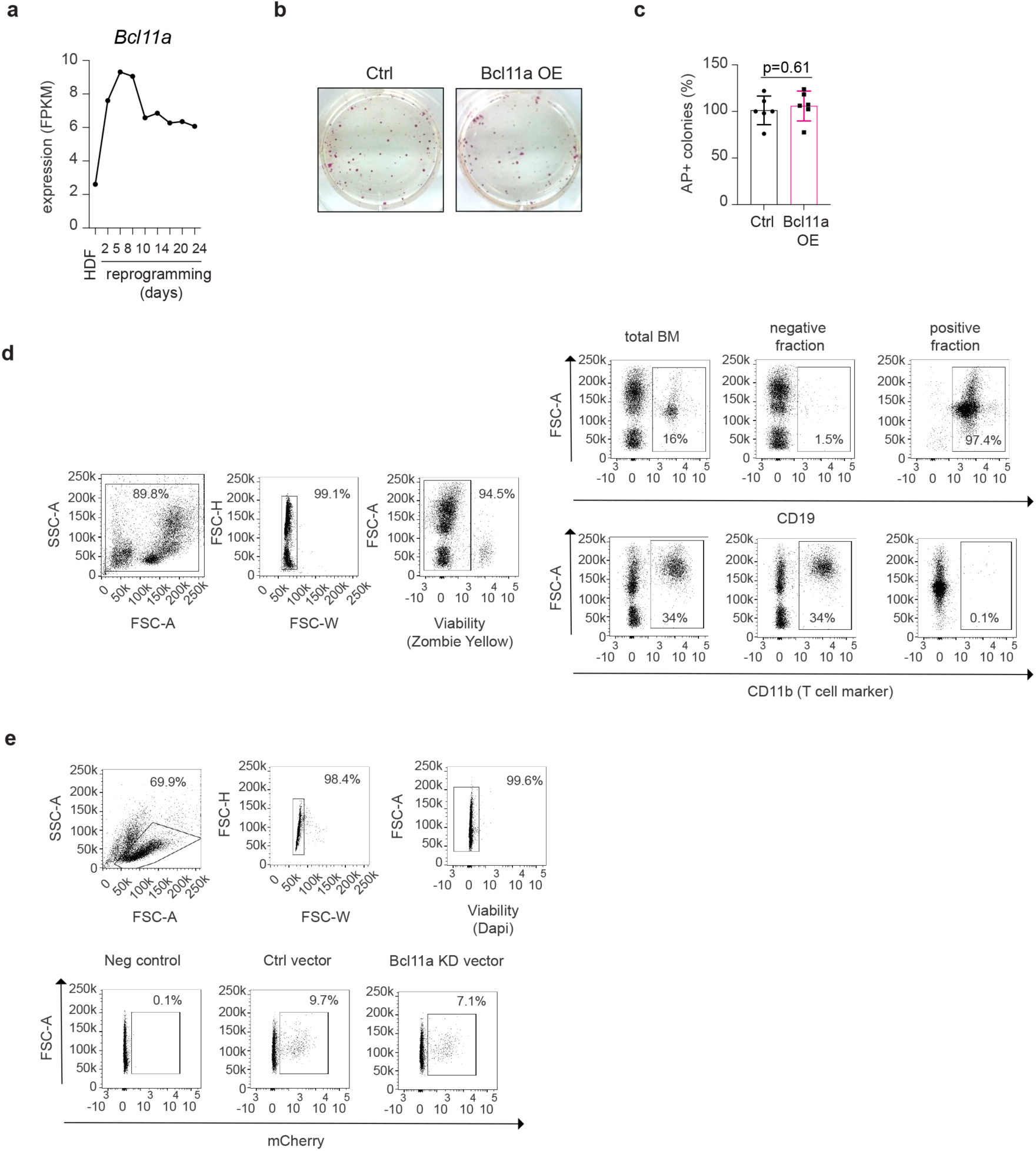
Bcl11a regulates the efficiency of reprogramming, transdifferentiation and transformation *in vitro*. **a** Level of expression of Bcl11a transcript during HDF reprogramming (extracted from ref^30^.). **b** Representative images of iPSC colonies stained for alkaline phosphatase (AP) activity upon Bcl11a OE. **c** Quantification of AP+ colonies (n=6 independent experiments). **d** FACS gating strategy for bone marrow (BM) cells (left panel) and subsequent analysis of expression of CD19 and CD11b in the total BM fraction as well as in the CD19-positive and CD19-negative fraction following CD19 positive MACS sorting (right panel). **e** FACS gating strategy for CD19-positive cells (upper panel) and analysis of mCherry expression in Ctrl and Bcl11a KD cells (lower panel). **f** Representative images of immortalized foci upon Bcl11a KD. **g** Quantification of foci (n=3 independent experiments). **h** Representative images of immortalized foci upon Bcl11a OE. **i** Quantification of foci (n=7 independent experiments). The data represent means ± sd (**c**, **g**, **i**). For statistical significance two-tailed student’s T-test (**c, g, i**).

**Supplementary Fig. 3:**
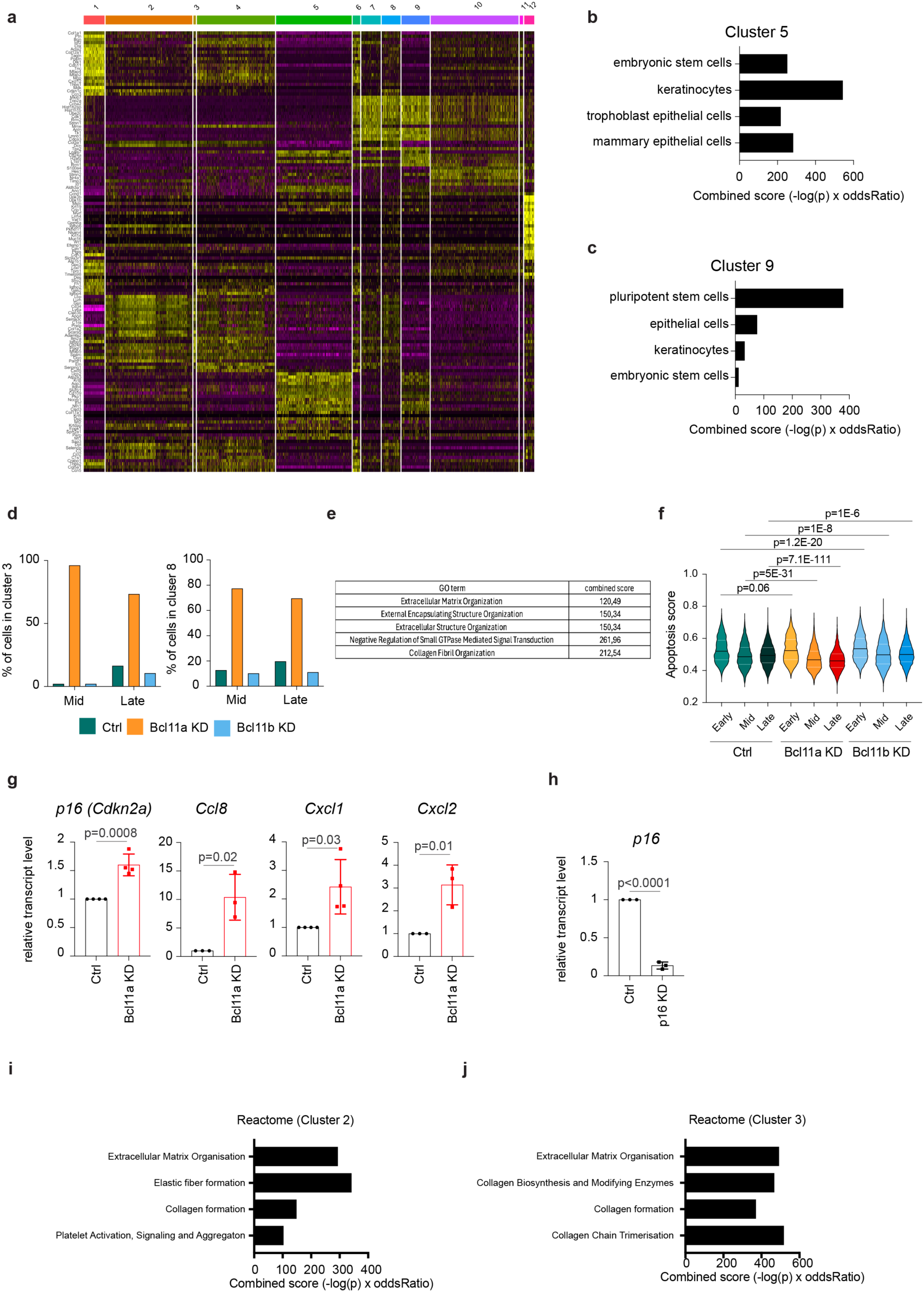
Depletion of Bcl11a and Bcl11b has different effects on reprogramming trajectories. **a** Heatmap of differentially expressed genes in the different clusters obtained from the single cell RNA-seq data. **b-c** Gene ontology (GO) terms associated with marker genes of Cluster 5 (**b**) and Cluster 9 (**c**) identified using the Panglao Database. **d** Proportions of cells assigned to each reprogramming stage across Cluster 3 (left) and Cluster 8 (right). **e** Gene ontology (GO) terms associated with marker genes of Cluster 4. **f** Level of apoptosis signature score from ref.^13^ at Early, Mid and Late reprogramming stages across samples. **g** RT-qPCR analysis of *p16, Ccl8, Cxcl1* and *Cxcl2* relative expression levels in Ctrl and Bcl11a KD condition at late reprogramming stage (n=3 or 4 independent experiments). **h** RT-qPCR analysis of *p16* expression in Ctrl and p16 KD MEF. (n=3 independent experiments). **i-j** GO terms associated with marker genes of Cluster 2 (**i**) and Cluster 3 (**j**) identified using the Reactome 2022 Database. The data in **(g, h)** are presented as means ± sd. Statistical significance was assessed using a two-tailed Student’s t-test (**g, h**) or by Wilcoxon Ranks Sum test (**f**).

**Supplementary Fig. 4:**
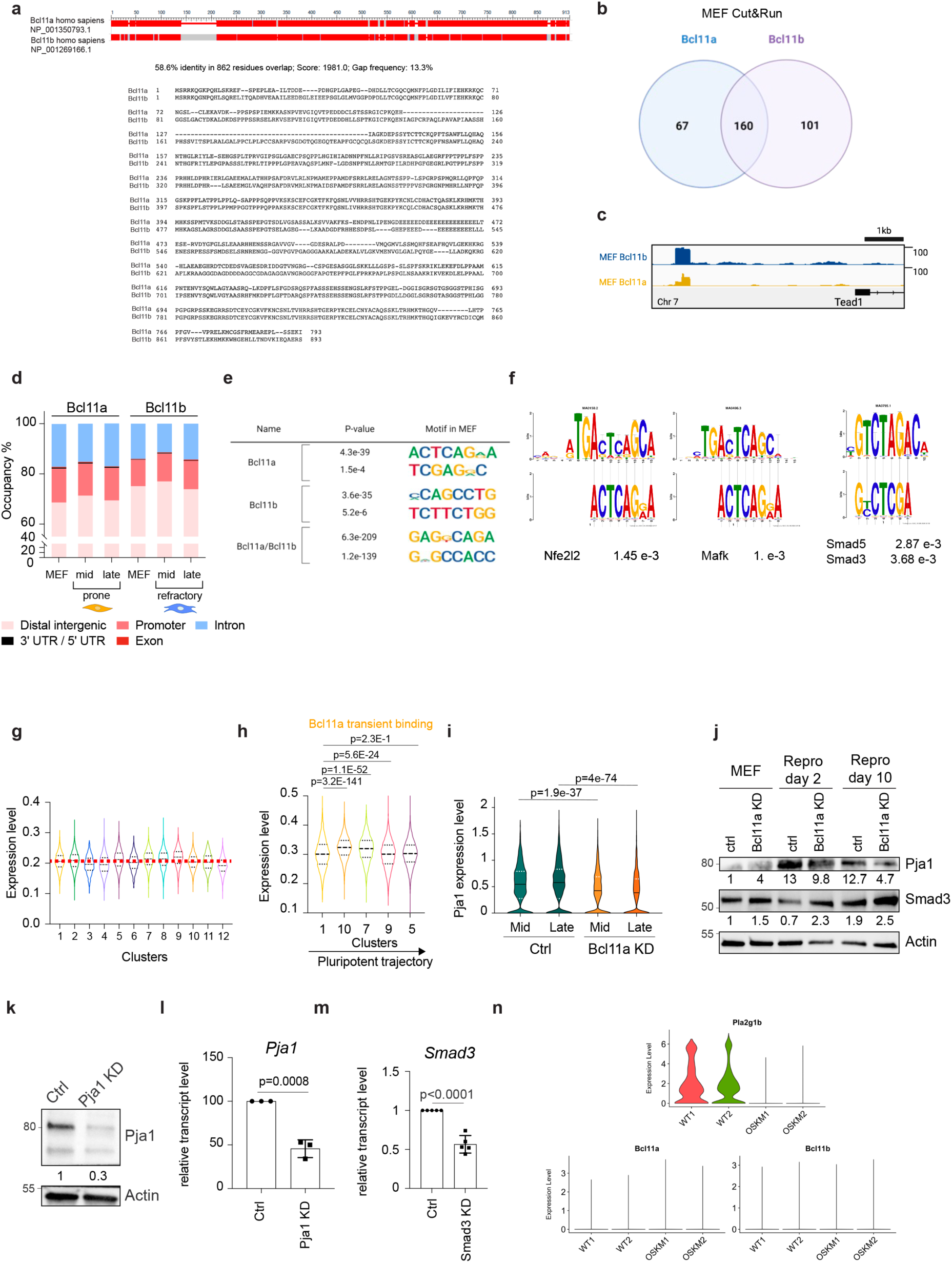
Bcl11a is redistributed by Oct4 during reprogramming. **a** Sequence homology between Bcl11a and Bcl11b genes. **b** Venn diagram of Bcl11a and Bcl11b bound genes in MEFs obtained by Cut&Run analysis. **c** Snapshot of indicated genomics data at the Tead1 genomic locus co-occupied by Bcl11a and Bcl11b. **d** Genomic distribution of Bcl11a and Bcl11b peaks. **e** Motif analysis by DREME of Bcl11a-specific, Bcl11b-specific and common binding sites in MEF. **f** Tomtom comparison of Bcl11a binding motif to Nfe2l2 (left panel) and Mafk (middle panel) and to Smad3/5 (right panel) in MEFs. **g** Graph depicts the mean expression level of the 89 genes that are bound by at least Bcl11a or Bcl11b throughout reprogramming. **h** Graph depicts the mean expression level of the 34 genes that acquired Bcl11a binding at the late stage of reprogramming. **i** Violin plot depicts *Pja1* expression level in Ctrl and Bcl11a KD settings at the Mid and Late stage of reprogramming. **j** Western blot of Pja1 and Smad3 levels in Ctrl and Bcl11a KD settings at early and late stage of reprogramming. **k** Western Blot for Pja1 and Actin in Ctrl and Pja1 KD MEF. **l** RT-qPCR analysis of *Pja1* relative expression level in Ctrl and Pja1 KD MEF (n=3 independent experiments). **m** RT-qPCR analysis of *Smad3* relative expression level in Ctrl and Smad3 KD MEF (n=5 independent experiments). **n** Violin plot depicting *Pla2g1b* (marker of somatic identity), *Bcl11a* and *Bcl11b* during *in vivo* reprogramming of pancreatic cells. Data are extracted from ref.^55^. The data in **l, m** are presented as means ± sd. Statistical significance was determined by a two-tailed Student’s t-test (**l, m**) or Wilcoxon Ranks Sum test (**h, i**).

## Methods

### Mice and MEFs

Animals were housed in a specific pathogen free (SPF) animal facility (P-PAC, Cancer Research Center of Lyon (CRCL), France). All experiments were performed in accordance with the healthcare regulations for animals used for scientific purposes governed by the European Directive 2010/63/EU. Protocols were validated by the local Animal Ethics Evaluation Committee (ACCeS: CSEA-10) and authorized by the French Ministry of Education and Research. R26^rtTA^; Col1a1^4F2A^ ^47^, Oct4-EGFP and Bcl11b-tdTomato mice were raised under standard conditions and bred in accordance with French national guidelines. Mice were genotyped using tail-derived genomic DNA. Tails were digested using DirectPCR Lysis reagent (102-T; Euromedex) and PCR was conducted using the Invitrogen Platinum II Hot-Start Green PCR Master mix (Life technologies).

MEF were extracted from mouse E13.5 embryos. The head and internal organs were removed and the embryo tissue was dissociated mechanically and then by trypsination for 10min at 37°C. Thus, cells were resuspended in MEF medium and propagated.

### Cell culture and viral production

MEFs were cultured in DMEM GlutaMAX medium supplemented with 10% of fetal bovine serum (FBS), 100 U ml^-1^ penicillin-streptomycin, 1 mM sodium pyruvate, 0.1 mM non-essential amino acids (NEAAs) and 0.1 mM b-mercaptoethanol. All MEFs used for the experiments were passaged maximum 4 times. CD19+ B cells from bone marrow were isolated with a CD19 (BD) antibody using MACS (Miltenyi Biotech) as previously reported^14^. B lymphocytes were cultured in RPMI medium supplemented with 10% of FBS, 100 U ml^-1^ penicillin-streptomycin, 0.1 mM b-mercaptoethanol and 10 ng ml^-1^ IL-7, IL-4, IL-15.

Retroviral particles were produced in Plat-E cells which were transfected in 10 cm dishes using CalPhos Mammalian Transfection kit (Ozyme) according to the manufacturer’s instructions. Following 7 h media were replaced with fresh MEF medium, which was collected after 48 h and the cells of interest were freshly transduced. Lentiviral vector particles were produced in 293FT cells using the same protocol. Additionally, during the transfection step the lentiviral vectors were co-transfected with plasmids encoding the G glycoprotein of the vesicular virus and Gag-Pol.

### Plasmids and constructs

PLKO, MSCV, Tet-O-FUW – Brn2, Ascl1, Myt1l, Sall4, Nanog, Esrrb, Lin28, FUdeltaGW-rtTA, pMXS-Oct3/4, pMXS-Sox2, pMXS-Klf4 and pMXS-cMyc plasmids were purchased from Addgene. Short hairpin RNA (shRNA) against Bcl11a, Bcl11b, Cdkn2a, p53, Smad3 and Pja1 were designed using the MERK Predesigned ShRNA interface and ligated into the AgeI and EcoRI digested pLKO.1 plasmid using the Rapid DNA ligation kit (Sigma-Aldrich). Overexpression vectors were obtained by amplifying the Bcl11a,Bcl11b and Pja1 genes from MEF cDNA by PCR and inserted into the BamHI digested MSCV vector using the In-Fusion HD Cloning Kit (Takara Bio). HA-Streptag-Turbo ID construction was added to the MSCV-Bcl11a OE vector by Vectorbuilder. PLV based vectors (PLV scramble/Bcl11a KD- mCherry) were obtained from Vectorbuilder. The pWPIR H-ras G12V and cyclin E plasmids were kindly supplied by the laboratory of A. Puisieux. For the luciferase assays, the Pja1 enhancer region was cloned into a CMV-luciferase vector by VectorBuilder company.

### Pluripotent reprogramming

R26^rtTA^; Col1a1^4F2A^, Oct4-EGFP or Bcl11b-tdTomato MEFs were plated at a density of 15,000 cells cm^-2^ (6-well plate) in MEF medium. The next day, cells were infected for 12 h (up to 48 h) with shRNA – carrying lentiviral stocks in the presence of 8 µg/mL of polybrene. Subsequently, the transduction medium was replaced with MEF medium supplemented with 2 µg/mL doxycycline. 48 h after doxycycline treatment, MEFs were reseeded at a density of 30,000 cells cm^-2^ (6-well plate) in 0.2% gelatin-coated dishes in iPSC medium (DMEM GlutaMAX medium supplemented with 15% of KnockOut Serum Replacement, 1000 U ml^-1^ leukaemia inhibitory factor, 100 U ml^-1^ penicillin-streptomycin, 1mM sodium pyruvate, 0.1 mM non-essential amino acids (NEAAs) and 0.1 mM b-mercaptoethanol). iPSC medium supplemented with doxycycline was changed every other day. Following 14 days of reprogramming AP staining was performed with the Leukocyte Alkaline Phosphatase Kit (Sigma-Aldrich) according to the manufacturer’s instructions.

### Oncogenic transformation

MEFs were infected overnight with shRNA carrying lentiviral stocks in the presence of 8 μg/mL polybrene. After 48 h, the cells were co-infected overnight with H-rasG12V, shTrp53- and Myc-carrying viruses. MEFs were reseeded 48 h post-infection in six-well plates at low density (500, 1,000 or 2,000 cells per well) in focus medium (MEF medium with 5% FBS) for the foci formation assay. The medium was then changed twice a week. After several passages of the cells derived from oncogenic transformation, soft agar assays were performed. Transformed cells were plated on an agarose-containing MEF medium layer at a density of 25,000-50,000 cells per six-well plate. Foci and soft agar colonies were stained 25-30 days later with a 0.5% cresyl violet solution in 20% methanol. shTrp53, H-rasG12V and cyclin E-expressing plasmids were also alternatively used in different combinations to induce oncogenic transformation.

### B cell reprogramming

CD19+ cells were extracted from OSKM Dox-inducible mice. They were then infected with PLV shRNA-carrying lentiviral stocks in presence of 10 µg/mL of protamine sulphate in B cell medium. Infected cells were FACS sorted using the mCherry reporter and plated at 20,000 cells cm^-2^ in gelatinized plates on mitomycin inactivated OP-9 feeder in B cell medium supplemented with 2 µg/mL doxycycline. After 48 h the medium was replaced with iPSC medium supplemented with doxycycline and 10 ng/mL of IL-7, IL-4 and IL-15. The medium was changed every 2 days.

### Xenografts

Suspensions of 100 µL of Matrigel and 100 µL containing 300 to 3.10^6^ transformed polyclonal cell lines were injected subcutaneously into female immunodeficient NMRI nude mice at the age of 10 weeks. 10 animals were used per group. Tumors were measured in size and volume every 3 days. Following a maximum of 16 days, tumors were extracted and fixed in 4% of paraformaldehyde for sectioning.

### Teratoma

Suspensions of 100 µL of PBS and 1.10^6^ iPSC were injected into the testis of 4 SCID mice at the age of 7 weeks. Following teratoma formation (4 weeks), tumors were extracted and fixed in 4% of paraformaldehyde for sectioning and hematoxylin-eosin staining.

### RNA extraction and real time quantitative PCR

RNA was isolated using Nucleospin RNA kit (Macherey-Nagel) and reverse transcribed using RevertAid H Minus First Strand cDNA Synthesis Kit (Life technologies). qPCRs were set up in triplicate using the ONEGreen FAST qPCR Premix (Macherey-Nagel) and primers (listed in supp table) on a Light Cycler 96 machine (Roche). Data were analyzed using the Light Cyler 96 1.1 software. Housekeeping genes used were Rs17 and Actin.

### Protein extraction and Western blot

Cell lysates were prepared in RIPA buffer (150 mM NaCL, 1% Triton, 0.5% deoxycholate, 0.1% SDS and 50 mM Tris at pH 8.0) with protease and phosphatase inhibitors. Proteins were extracted as previously reported^4^. Proteins were separated on SDS-PAGE gels and subsequently transferred to nitrocellulose membranes. Membranes were blocked using 5% of milk in TBS-Tween for 1 h and probed with a primary antibody overnight at 4°C. After washing, the membranes were incubated with the HRP-coupled secondary antibody 1 h at room temperature. Antibody signal was developed using ECL and DURA reagents. Data were obtained and quantified in Bio-Rad Image Lab Software.

### Cut&run

Bcl11b-tdTomato reprogrammable MEFs were expanded and reprogrammed to pluripotency. Every 3 days reprogramming samples were stopped and cells were FACS-sorted for the expression of Bcl11b and Thy1.

Cut and run experiments were carried out as previously described with some modifications^56^. Briefly, 250,000 cells were captured with BioMagPlus Concanavalin A (Polysciences, 86057-3) and incubated with the appropriate primary antibody overnight at 4°C. For studying the TF *Bcl11a*, a combination of two antibodies was used (Bethyl Laboratories, A700-073, 3 µg/100 µL and Abcam, ab191401, 3 µg/100 µL). For the *Bcl11b* TF, another combination was used (Bethyl Laboratories, A300-385A, 3 µg/100 µL and Abcam, ab240636, 2 µg/100 µL). The following day, the samples were incubated with a secondary antibody at room temperature for 1 h (Donkey anti-rabbit IgG antibody: Sigma-Aldrich, SAB3700932; 2 µg/100 µL). After washing, pAG-MNase enzyme (Cell Signaling, 57813, dilution 1:33) was added and incubated for 1 h at 4°C, then activated by adding CaCl_2_ (2 mM final concentration) for 30 min. After stopping the reaction, the fragmented chromatin was released by centrifugation and digested with proteinase K (20 mg/mL) (Life technologies 25530049) overnight at 37°C. The next day, DNA was extracted by phenol/chloroform (Sigma-Aldrich 77617) followed by ethanol precipitation.

Cut and Run DNA libraries were prepared using the NEBNext Ultra II DNA Library Prep Kit (New England BioLabs, NEB E7645S) according to the manufacturer’s instructions. For PCR amplification, the primers were used at a concentration of 0.2 µM with the following cycling conditions: 98°C for 30 s, 15-20 cycles of 98°C for 10 s and 65°C for 75 s; followed by a final extension at 65°C for 5 min. DNA libraries were cleaned using 1x volume of SPRIselect beads (Beckman Coulter, B23318) and Illumina sequenced (150-nt pair-end sequencing; Illumina NovaSeq 6000).

### Immunoprecipitation Mass Spectrometry (IP-MS)

R26rtTA; Col1a14F2A MEFs were infected with retroviral particles carrying empty vectors or vectors overexpressing Bcl11a-streptag or Bcl11b-streptag. After 72 h, MEFs were lysed in IP lysis Buffer (20 mM Tris pH 7.5, 150 mM NaCl, 1% NP40) supplemented with proteases and phosphatases inhibitors. Immunoprecipitation assays were conducted using Strep-tag II beads (IBA MagStrep XT beads). 1.5-2 mg of protein lysate was incubated with beads overnight by rotating on a wheel at 4°C. The following day, the immunocomplexes were washed three times with washing buffer (20 mM Tris pH 7.5, 150 mM NaCl) using a magnetic rack. Proteins were eluted in 2x Laemmli buffer (40 mM Tris HCL pH 6.8, 2% SDS, 20% glycerol and 0.02% bromophenol blue) at 95°C for 10 min. IP eluates were Silver stained using SilverQuest™ Silver Coloring Kit (Thermofischer).

IP beads were prepared with the iST kit (ref. P.O.0001, PreOmics) according to the manufacturer’s protocol for Magnetic Immunoprecipitation Samples. Briefly, proteins were denatured, reduced and alkylated (10 min, 60°C, 1,000 rpm), then digested with a mixture of endoproteinases Lys-C/trypsin (3 h, 37°C, 500 rpm). After a cleaning step, peptides were dried, suspended in 50 µL Formic Acid (FA) 0.1%.

1 µL of each sample was analyzed on the Exploris 480 mass spectrometer coupled with a Vanquish NEO nanoLC system (ThermoFisher Scientific). Peptide samples were loaded onto a C18 Acclaim PepMap100 trap-column 300 µm ID x 5 mm, 5 µm, 100Å (ThermoFisher Scientific) and separated on a C18 Acclaim Pepmap100 nano-column, 50 cm x 75 µm i.d, 2 µm, 100 Å (ThermoFisher Scientific) with a 30 min linear gradient from 3% to 35% buffer B (A: 0.1% FA in H2O, B: 0.1% FA in ACN/H2O-80/20-v/v) followed by 10 min at 100% B and then by a washing and equilibration steps for a total duration of 42 min. The flow rate was 300 nL/min and the oven temperature was kept constant at 45°C. Peptides were analyzed with a DDA 1s HCD method: MS data were acquired in a data-dependent strategy (DDA) selecting the fragmentation events based on the most abundant precursor ions in a 1s survey scan (350-1400 Th). Resolutions of the survey and MS/MS scans were respectively set at 120,000 and 15,000 at m/z 200 Th. The Ion Target Values for the survey and the MS/MS scans in the Orbitrap were set at 3E6 (300%) and 1E5 (100%), respectively, and the maximum injection time was set at 50 ms for MS scan and 22 ms for MS/MS scan. Parameters for acquiring HCD MS/MS spectra were as follows: collision energy = 30 and isolation window = 4 m/z. The precursors with unknown charge state, charge state of 1 and 8 or greater than 8 were excluded. Peptides selected for MS/MS acquisition were then placed on an exclusion list for 40 s using the dynamic exclusion mode to limit duplicate spectra.

### FACS analysis

Oct4-GFP, Bcl11b-tdTomato and other gene expression were analyzed using antibodies (listed hereafter) on a BD LSRFortessa with the FACS Diva version (version 8.0). Data were analyzed using FlowJo version 10 software.

### MEFs to neurons transdifferentiation

Wild-type MEFs were co-infected with FUdeltaGW-rtTA and shRNA (control or targeting Bcl11a) lentiviral plasmids 2 days before the experiment in the presence of 8 μg/mL polybrene. At day 0, the cells were co-infected with Tet-O-FUW-Brn2, -Ascl1 and -Myt1l lentiviral plasmids. The following day, the medium was replaced with fresh MEF medium supplemented with 2 μg/mL doxycycline. At day 3, the medium was replaced with fresh N3 medium consisting of DMEM/F12, 100 U ml−1 penicillin/streptomycin, 2.5 μg/mL insulin, 50 μg/mL apo-transferrin, 86.5 μg/mL sodium selenite, 6.4 ng/mL progesterone and 16 μg/mL putrescine supplemented with 2 μg/mL doxycycline. The medium was changed daily until day 7–8.

### Immunofluorescence

Cells were fixed in 4% paraformaldehyde for 10 min at room temperature, washed three times with PBS, permeabilized with 0.1% Triton X-100 for 30 min at room temperature and blocked with 1% bovine serum albumin for 1 h. After incubation with primary antibody overnight at 4°C, the cells were washed three times with PBS and incubated with fluorophore-labelled appropriate secondary antibody (Life Technologies). Acquisition was done with Axiovision 4.8.2 software.

### Luciferase assay

HEK293 cells (100.000 cells/well in 24-well plates) were transfected using 400ng reporter vector (Firefly Luciferase), 5ng pRL-SV (Renilla Luciferase control vector, and 300ng expression vectors (when necessary). Cell lysates were assayed for luciferase activity two days post-transfection using the Promega Dual Luciferase kit and a Spark Tecan microplate luminometer. Firefly Luciferase activity was reported to Renilla Luciferase activity to normalize for the differences in transfection efficiencies. Each experiment was repeated three times, and each measurement was performed in duplicates.

### ScRNA-seq

All analyses were conducted using R (version 4.3.2) and the Seurat package (version 5.0.1)^48^. All visualizations were performed using the dyno package in R (ref: https://github.com/dynverse/dyno). Single-cell RNA sequencing (scRNA-seq) data for both the Reprogramming-Transformation-Transdifferentiation (ReproTransfoTransdiff) and the MEF reprogramming assays were prepared, dissociated, and sequenced following the protocol outlined in our previous publication^4^, except that the 3’ assay of 10x Chromium technology (v3 chemistry) was used, with all samples multiplexed together in each assay.

### Data quality control (QC) and filtering

For the ReproTransfoTransdiff dataset, cells were filtered based on mitochondrial content and gene expression. Cells with more than 15% mitochondrial reads were removed. Basic quality metrics such as the number of detected genes, total reads per cell, and the ratio of mitochondrial reads were examined using violin plots generated with Seurat. A distinct bimodal distribution was observed in the number of detected genes per cell, and cells representing the lower mode of the bimodal distribution were excluded by setting a threshold of 3,000 genes per cell minimum. Additionally, MEFs expressing mean logcounts of Thy1 and Bcl11b below 1 were filtered out. After filtering, the final dataset contained 1,838 MEFs, 1,245 transforming cells, 2,633 reprogramming cells and 869 transdifferentiating cells. Counts were normalized and scaled with the number of counts per cell and mitochondrial read ratio included as regression variables.

For the MEF reprogramming in the KD condition dataset, the number of cells per sample ranged from 19,401 to 23,757. Similarily to the ReproTransfoTransdiff dataset, cells pertaining to the lower mode of bimodal distributions in gene detection and read counts, likely representing damaged or dying cells, were filtered out by setting a minimum threshold of 4,000 genes per cell. Additionally, cells with more than 10% mitochondrial reads were removed, leaving between 4,752 and 5,863 cells per sample after filtering.

### Low-dimensional embeddings

A diffusion map for the ReproTransfoTransdiff dataset was generated using the destiny R package (version 3.16.0)^49^, based on the first two principal components of the MNN-integrated embedding, calculated using fastMNNIntegration function from the SeuratWrappers package (ref 1: https://github.com/satijalab/seurat-wrappers) with default settings. For the MEF reprogramming in KD conditions dataset, Uniform Manifold Approximation and Projection (UMAP) dimensionality reduction was performed using Seurat’s RunUMAP function with default parameters, utilizing the first 30 dimensions of the PCA embedding.

### Clustering and marker gene calculation

For both the ReproTransfoTransdiff and the MEF reprogramming in KD condition datasets, we employed density-based clustering using the dbscan R package^51^ on the 10-dimensional UMAP embedding, which was specifically computed only for that purpose using Seurat’s RunUMAP function on the first 30 PCA components (with n.neighbors = 15 and min.dist = 0). Clusters were assigned using the dbscan::optics function with minPts set to 100 and eps_cl set to 0.5. This approach identified 5 and 18 clusters, respectively, although approximately 5% in the ReproTransfoTransdiff and 10% of cells in the KD dataset could not be assigned to any cluster and were excluded. Additionally, in the case of the KD dataset, six clusters containing fewer than 200 cells were also removed.

### Trajectory inference

Cellular trajectories were inferred using the slingshot R package, based on the 2-dimensional UMAP embedding and the dbscan-derived clusters as the clusterLabels parameter. Cluster 1 was designated as the starting point for trajectory analysis.

### Cut&run data analysis

Peak calling was performed using MACS2, and the resulting BED files were imported into R (v4.3.2) for further analysis. ChIPseeker (v1.38.0, 10.18129/B9.bioc.ChIPseeker) and ChIPpeakAnno (v3.36.1, 10.18129/B9.bioc.ChIPpeakAnno) packages were employed to annotate and explore the peaks. Binding site overlaps between samples were identified using Bedtools (v2.3.0)^52^. To investigate chromatin states, the peaks from our CUT&RUN data were crossed with data from Chronis et al.,2017 using Bedtools intersect function. The intersected peaks were binarized using the BinarizeBed function from the ChromHMM suite (v1.25) and then analyzed using the LearnModel function within the same suite. Motif analysis was carried out using the memes R package (v1.3.2) (https://snystrom.github.io/memes/), which serves as a wrapper for the MEME Suite^53^. BedGraph files generated from the peak caller were visualized using IGV^54^ for manual inspection and visualization of peak distributions.

### IP-MS data analysis

Raw data were processed with Proteome Discoverer 3.1 through the SEQUEST HT and CHIMERYS search engines against the uniprot Mus musculus database (Release november 2023, 88384 sequences) and a database of common contaminants. Precursor mass tolerance was set at 10 ppm and fragment mass tolerance at 0.02 Da, and up to 2 missed cleavages were allowed. Oxidation (M), acetylation (Protein N-terminus) were set as variable modification and Carbamidomethylation (C) as fixed modification. Validation of identified peptides and proteins was done using a target decoy approach with a false discovery rate positive FDR < 1%.

### Data availability

The next generation sequencing data of the study are deposited in GEO omnibus under the accession number GSE283626 (reviewer token qbarusiqrzyljud).

## Acknowledgements

This work was supported by the Fondation Bettencourt Schueller (Impulscience 2022 to FL), La Ligue Contre le Cancer Nationale et Régionale (to F.L.), Institut National du Cancer (2019-L22 to F.L.), Agence Nationale de la Recherche (Stemnet to F.L), Fondation pour la recherche médicale (to A.T.), Fondation ARC (to A.H.), Centre Léon Bérard (F.L.), Labex DevWeCan (ANR-10-LABX-0061 to F.L), Institut Convergence PLAsCAN (ANR-17-CONV-0002). We thank the core facilities of the Centre de Recherche en Cancérologie de Lyon (CRCL) and Centre Léon Bérard (CLB), and in particular Cyril Degletagne (Cancer Genomics Platform, CGP, CRCL) for single cell RNA-seq and Thibault Andrieu (Flow Cytometry Core Facility, CYLE, CRCL) for FACS analysis. We acknowledge Frédéric Delolme and Adeline Page from Protein Science Facility of SFR Biosciences (UAR3444/CNRS, US8/Inserm, ENS de Lyon, UCBL) for Mass Spectrometry analysis. We thank Brigitte Manship for proofreading the manuscript. BioRender was used for illustrations.

## Author contributions

A.T. performed most of the experiments presented in the figures. M.R., I.K, I.M, L.C. and B.M. contributed to the generation of data. E.Z. and Y. T. performed the bioinformatics analyses. M.F. and M.M.P. conducted Cut&Run experiments related to Figure 6. N.G. and F.F. conducted the histologic analyses of teratoma. E. H., C. H., N.M., D. B., E. D contributed to generate data that were not incorporated in the manuscript. Y. G. B. contributed to the design and to the conduction of B-lymphocytes reprogramming presented in Figure 2. A.T., A.H. and F.L designed experiments for the manuscript. A.T., A.H., E.Z. and F.L wrote the manuscript. A.H and F.L. designed and supervised the study. All authors approved and contributed to the final version of the manuscript.

## Materials and Correspondence

Correspondence and material requests should be addressed to A.H. and F.L.

## Supplementary information

Supplementary Figures 1-5 and Supplementary Table 1.

